# Glycosylceramide assembly and function in a model bryophyte

**DOI:** 10.64898/2026.07.31.742004

**Authors:** Pauline Duminil, Sönke Beewen, Merle Reinhold, Deborah Oliveira Schopp, Stefanie König, Cornelia Herrfurth, Ivo Feussner, Tegan M. Haslam

## Abstract

**Research Conducted:** To elucidate the functions of glycosylceramides, we generated and characterized mutants deficient in multiple steps contributing to their assembly in the model moss *Physcomitrium patens*. We mutagenized *SPHINGOLIPID Δ8-DESATURASE*, whose products are preferentially incorporated into glycosylceramides, and a suite of higher-order mutants combining *sphingolipid Δ4-desaturase* and *glycosyl ceramide synthase*.

**Methods:** We used targeted lipidomics to describe the chemotypes of all mutants. We used quantitative phenotype analysis, transcriptomics, and phytohormone profiling to understand the effects of these chemotypes on development and physiology.

**Key Results:** These mutants present a range of phenotypes that collectively indicate that in *P. patens* (1) glycosylceramide deficiency impairs development, largely due to imbalance in free ceramide homeostasis (2) the synthesis of glycosylceramides is dependent upon the presence of a specific free ceramide profile (3) the Δ4-, but not the Δ8-desaturation, is strictly required for glycosylceramide synthesis, (4) cell division and differentiation, but not cell expansion, are affected by sphingolipid imbalance, and (5) sphingolipid imbalance results in oxylipin accumulation.

**Conclusion:** Collectively, our results elucidate the assembly and functions of glycosylceramides in a model bryophyte, and highlight conserved and specialized aspects of sphingolipid metabolism among plants.

## Introduction

Sphingolipids are ubiquitous and essential membrane components in eukaryotic cells. Two classes of complex sphingolipids can be distinguished based on the linkage between their lipidic ceramide backbone and polar headgroup: Complex phosphosphingolipids have a phosphodiester linkage, whereas complex glycosphingolipids have a glycosidic bond (Ternes *et al*., 2011). Complex phosphosphingolipids are present in plants as glycosyl inositol phosphorylceramides (GIPCs). These have a large headgroup with a core glucuronic acid-inositol phosphate structure, which is usually further glycosylated (reviewed in (Mortimer & Scheller, 2020)). GIPCs make up approximately one quarter of the plasma membrane lipidome (Bahammou *et al*., 2024), and contribute to its thickness (Levine *et al*., 2000) and order (Lenarčič *et al*., 2017). They have defined roles in plant-microbe interactions (Lenarčič *et al*., 2017; Moore *et al*., 2021; Mittendorf *et al*., 2025), salt sensing (Jiang *et al*., 2019), polar membrane sorting (Ito *et al*., 2021; Mittendorf *et al*., 2025), and cell division (Molino *et al*., 2014; Wegner *et al*., 2025). Glycosphingolipids are present in plants as glycosylceramides (GlcCer, sometimes also referred to as HexCer, or GluCer). These usually have a single monosaccharide as their headgroup, and the sugar has often been identified as glucose (reviewed in (Haslam & Feussner, 2022)). GlcCers are estimated to make up about one third of the total sphingolipid pool in *Arabidopsis thaliana* (Markham *et al*., 2006), yet relatively little is known about their *in planta* functions.

GlcCers and the core ceramide structures they are built from are synthesized in the endoplasmic reticulum; beyond this, the precise subcellular distribution of GlcCers is unclear, but they have been found enriched in tonoplast membranes in Arabidopsis, *Hordeum vulgare*, *Kalanchoë daigoremontiana*, and *Mesembryanthenum crystallinum* (ice plant)(Haschke *et al*., 1990; Carmona-Salazar *et al*., 2021; Guo *et al*., 2022). Mutant characterization in *A. thaliana* revealed a seedling-lethal phenotype when the committed step of GlcCer assembly, headgroup addition by GLYCOSYLCERAMIDE SYNTHASE, was knocked out (Msanne *et al*., 2015). *In vitro* cultivation of the *Atgcs* mutant as callus culture revealed a defect in organogenesis, suggesting a role for GlcCer in differentiation (Msanne *et al*., 2015). Transmission electron microscopy of *gcs* mutants showed that individual Golgi stacks contained significantly fewer cisternae (Msanne *et al*., 2015). In contrast to *gcs* mutants, RNAi down-regulation of *AtGCS* produced viable plants, even though they presented up to 98 % depletion of total GlcCer. The RNAi lines were severely stunted, but could complete vegetative and reproductive development. This suggests that a very small amount of GlcCer is sufficient to sustain growth, albeit with growth defects (Msanne *et al*., 2015).

For further insight into the molecular functions of GlcCer, we previously generated and characterized knock-out *gcs* mutants of the model moss, *Physcomitrium patens* (Gömann *et al*., 2021a). Knock-out *Ppgcs* mutants had <1 % of wild-type GlcCer levels, and accumulated free ceramide precursors of GlcCer. They also hyperaccumulated GIPCs. Perhaps surprisingly, *Ppgcs* mutants were viable, although they had a stunted phenotype—analogous to *GCS* RNAi lines of *A. thaliana*.

Profiling the lipidome of *P. patens* had already revealed that its GlcCer pool is composed almost exclusively of a single ceramide backbone, 18:2;2/20:0;1 (Resemann *et al*., 2021). (Standard sphingolipid nomenclature describes first the long chain base (LCB) moiety, followed by (/) the fatty acyl moiety. The three numbers included for each indicate the chain length (:) number of desaturations (;) and number of hydroxylations). In *P. patens*, the 18:2;2/20:0;1 ceramide is not a major component of the free ceramide pool, and it is not incorporated into GIPCs. Similar specific prevalence of an 18:2;2 LCB moiety in GlcCer is common, but not ubiquitous, across land plants (Markham *et al*., 2006). Of the ceramide modifications required to synthesize 18:2;2/20:0;1, the Δ4-desaturation of the LCB catalyzed by SPHINGOLIPID Δ4-DESATURASE (SD4D) was of particular interest. In plants such as *P. patens* where Δ4-desaturation is detected, its preferential incorporation into GlcCer appears consistent (Markham *et al*., 2006). In contrast, in *A. thaliana*, Δ4-desaturation is exclusively detected in flowers, and there at very low levels (approx. 4 % of GlcCer)(Michaelson *et al*., 2009). Loss of SD4D by mutagenesis in *A. thaliana* had no obvious effect on growth and physiology.

We questioned the metabolic and physiological significance of the Δ4 desaturation in *P. patens* (and in other plants with high SD4D activity) and therefore generated *P. patens sphingolipid Δ4-desaturase* (*Ppsd4d*) mutants (Gömann *et al*., 2021a). We found that *Ppsd4d* knock-out mutants lacked Δ4-desaturated sphingolipids and had severely depleted GlcCer, at less than 1 % of wild-type levels—though slightly more than the trace amount detected in *Ppgcs*. This showed that LCB Δ4-desaturation is not only preferred, but essential for flux into GlcCer in *P. patens*. Surprisingly, unlike *Ppgcs*, *Ppsd4d* mutants appeared nearly wild type-like (Gömann *et al*., 2021a).

Our characterization of *Ppgcs* and *Ppsd4d* raised several questions. First, why is the *Ppgcs* phenotype so much more severe than *Ppsd4d*? Is the observed slightly-higher residual amount of GlcCer present in *Ppsd4d* sufficient to sustain normal growth, or is the free ceramide profile that accumulates in *Ppgcs*, enriched in the Δ4-desaturated products of SD4D, among other substantial shifts, causing its severe phenotype? Second, having observed that Δ4-desaturation is critical for ceramide flux into GlcCers, what is the role of LCB desaturation at the Δ8 position, which is also strongly, preferentially incorporated into the 18:2;2/20:0;1 GlcCers of *P. patens*? Third, why does the *Ppgcs* mutant have such a severe morphological phenotype? To answer these questions, we expanded our collection of *P. patens* mutants: We generated *gcs sd4d* double mutants to determine whether suppressing the accumulation of free ceramide precursors in the *gcs* background could restore a normal phenotype, and *sd8d* and *sd4d sd8d* mutants to determine the effect of Δ8-desaturation on ceramide channeling into GlcCers. Then, moving beyond on the chemotypes of the mutants, we applied a suite of analyses for anatomical and morphological phenotype characterization. Based on the *gcs*, *sd4d*, *sd8d*, *sd4d gcs*, and *sd4d sd8d* mutants, we characterize the metabolism and functions of GlcCer in *P. patens*.

## Materials and methods

### *P. patens* cultivation

#### Standard conditions

Conditions used unless otherwise indicated were 16 h light/8 h dark, 105-120 μmol m^-2^ s^-1^, at 25 °C/18 °C. Protonema was grown on BCD-AT medium (1 mM MgSO_4_, 1.84 mM KH_2_PO_4_, 10 mM KNO_3_, 45 μM FeSO_4_, 5 mM ammonium tartrate, 1 mM CaCl_2_, Hoagland’s trace elements, 0.55 % plant agar) overlaid with sterile cellophane discs, and cultivated every seven to ten days by dispersing with an ULTRA-TURRAX (IKA) in sterile tap water. Gametophores were grown on BCD medium (BCD-AT medium lacking ammonium tartrate) (Maronova & Kalyna, 2016), and cultivated every five weeks by transferring pieces of material to fresh plates.

#### Cold stress

To exacerbate the *gcs* mutant phenotype, plates containing five-week-old gametophores grown in normal conditions were transferred and grown for six more weeks in autumnal conditions: short days (8 h light/16 h dark) at a constant 17 °C. Harvested material was frozen in liquid nitrogen and lyophilized for phytohormone and RNA extractions.

#### Growth assays

For gametophore growth assays, 1 mm protonema spot inocula were cultivated on BCD medium, with all genotypes for comparison cultivated on the same plate, equidistant from the center. For skotonema cultivation, 2 to 3 mm protonema spot inocula were placed on square plates with BCD-AT medium supplemented with 2 % (w/v) sucrose (Saavedra *et al*., 2015). These were grown in standard conditions for one week, then wrapped in parafilm, transferred to a dark growth chamber at 25 °C, and grown vertically for four weeks. For rhizoid growth assays, solid BCD medium was removed from one half of a square plate, then gametophores were transferred upright on the cut surface of the medium (Fu *et al*., 2022). The plates were placed vertically, allowing for visualization of rhizoid growth into the agar.

### Mutagenesis in *P. patens*

#### Guide design and mutagenesis

*Physcomitrium patens* ecotype Gransden was obtained from the International Moss Stock Center (IMSC; https://www.moss-stock-center.org/en/, strain #40001), used as a reference in all experiments, and as a genetic background for mutagenesis. The gene loci investigated are summarized in Table 1. The single *sd4d* mutant used here was generated by homologous recombination and was described previously as *sd4d-1*; the single *gcs* mutant used here was previously generated by CRISPR/Cas9 mutagenesis and described previously as *gcs-3* (Gömann *et al*., 2021a). *PpSD8D* was identified on Phytozome (https://phytozome-next.jgi.doe.gov/) where it is annotated according to sequence and domain similarities. CRISPR/Cas9 was used to generate (1) *sd8d* single mutants in the wild-type background, (2) *sd4d gcs* double mutants by mutagenizing *SD4D* in the *gcs* background, and (3) *sd4d sd8d* double mutants by simultaneously targeting both genes in a wild-type background. All of the mutant lesions are specified in Supplemental Table S1.

**Table 1:** Gene loci and annotations.

| Table 1: Gene loci and annotations |  |  |
| --- | --- | --- |
| Abbreviation | Genome annotation V3.3 | Genome annotation V6 |
| GCS | Pp3c16_7990V3.1 | Pp6c16_5700V6.1 |
| SD8D | Pp3c25_1720V3.1 | Pp6c25_6120V6.1 |
| SD4D | Pp3c23_19370V3.1 | Pp6c23_9600V6.1 |

CRISPR-Cas9 mutants were generated as described previously (Lopez-Obando *et al*., 2016; Collonnier *et al*., 2017; Haslam *et al*., 2024). Target sites were selected using the CRISPOR website (http://crispor.tefor.net/) and synthesized as oligonucleotides (Supplemental Table S2) to be cloned in the pUCRISPR vector. The plasmid was validated by colony PCR and sequencing. The other plasmids required for gene editing by CRISPR-Cas9, Act1pro-Cas9 and pBNRF, were provided by Prof. Fabien Nogué (INRA Versailles).

#### Protoplast transformation and mutant identification

Protoplast transformation methods were adapted from (Liu & Vidali, 2011; Maronova & Kalyna, 2016). The protoplasts from *P. patens* were generated using approximately six-day-old protonema incubated with Driselase (1 % w/v) and 0.44 M mannitol (8 % w/v) for three hours. All steps were carried out at RT. The digested cells were filtered through a sterile 70 µm mesh, then centrifuged at 130 rcf for 5 min and washed twice with 15 mL 0.44 M mannitol solution. The cells were then resuspended in MMg solution (0.4 M mannitol, 15 mM MgCl_2_, 4 mM MES pH 5.7). After incubation at room temperature for 30 min, 600 µL aliquots of the cells were mixed with equal amounts of guide plasmids and the pBNRF selection plasmid (approx. 5 µg each), 10 µg of pAct1-Cas9, and 700 µL of PEG-calcium solution (4 g PEG4000, 3 mL H_2_O, 2.5 mL 0.8 M mannitol, 1 mL 1 M CaCl_2_). The cells were incubated at room temperature for another 30 min. Finally, cells were washed with 3 mL W5 solution (154 mM NaCl, 125 CaCl_2_, 5 mM KCl, 2 mM MES pH 5.7) and centrifuged at 130 rcf for 5 min before resuspension in 5 mL molten, 42 °C, PRM-T medium (BCD-AT with 10 mM CaCl_2_, 0.33 M mannitol, 0.4 % (w/v) plant agar) to be spread over two plates containing PRM-B medium (BCD-AT with 10 mM CaCl_2_, 0.33 M mannitol) and cellophane.

After one week, cellophane discs containing recovered protoplasts were transferred to BCD-AT plates containing G418 for selection of resistant lines. Approximately two weeks later, resistant plants were transferred to fresh BCD plates. After enough material had grown, a piece of each resistant plant was sampled to extract genomic DNA (Extract-N-Amp™ Plant Tissue PCR Kits, Merck) and genotyped by amplifying and sequencing the targeted region (primers in Supplemental Table S2).

### RNA extraction for cloning and RNAseq

For cloning, total RNA was extracted from 5 mg lyophilized protonema using an E.Z.N.A plant RNA Kit (Omega Bio-Tek) according to the manufacturer’s instructions. The extract was treated with DNAse (Thermo Fisher Scientific), the RNA concentration was determined with a Nanodrop Spectrophotometer, and 1 µg was used to synthesize cDNA with the RevertAid H Minus Reverse Transcriptase (Thermo Fisher Scientific).

For RNA sequencing, total RNA was extracted from 10 mg lyophilized material using the Spectrum™ Plant Total RNA Kit (Sigma) with minor modifications for the use of lyophilized, rather than fresh, material. RNA concentration and quality were controlled via Nanodrop Spectrophotometer and an agarose gel. 1 µg of each sample was sequenced by Novogene (UK). WT control and stress treatments were deposited at https://rshiny.gwdg.de/apps/streptonet/, described in (Dadras *et al*., 2025). RNAseq data was analyzed by Novogene, and additional analyses presented here were also performed using R programming language (v4.5.3) libraries (R Core Team, 2026). DESeq2 (Love *et al*., 2014) was used for differential gene expression analysis. For DEG analysis raw counts were pre-filtered to retain only transcripts with ≥10 counts in ≥3 samples. ClusterProfiler (Xu *et al*., 2024) was used for GO and Pfam enrichment against GO.db (Carlson *et al*.), org.At.tair.db (Carlson, 2026) and InterPro REST API (Nightingale *et al*., 2017) as libraries. ggplot2 (Wickham, 2016) and ggrepel (Slowikowski, 2026) were used for visualization.

### Heterologous expression of *PpSD8D* in *Komagaetella phaffii delta8Δ* mutant

The *PpSD8D* coding sequence was amplified using the PfuUltra HF DNA polymerase (Agilent) for direct cloning in the pIB4 vector (Sears *et al*., 1998) (cloning primers are listed in Supplemental Table S2). The *Komagaetella phaffii* (formerly *Pichia pastoris*) GS115 WT strain (Invitrogen) and *delta8Δ* mutant (Ternes *et al*., 2011) were used as controls. Competent cells of the *delta8Δ* mutant were generated and transformed using the Pichia EasyComp kit (Thermo Fisher Scientific). Transformed cells were selected on Regeneration Dextrose Medium lacking histidine (1 M sorbitol, 1 % (w/v) dextrose 1.34 % (w/v) yeast nitrogen base, 4*10^-5^ % (w/v) biotin and 0.005 % (w/v) amino-acids) and verified by colony PCR (Supplemental Table S2) and four independent colonies were retained. The cultivation of *K. phaffii* was done according to (Weidner *et al*., 2010; Mohammadzadeh *et al*., 2021) with minor modifications. A 4 mL overnight pre-culture in Buffered Glycerol-complex Medium (BMGY) (Weidner *et al*., 2010) was prepared for each replicate. The next day 1 OD_600_ was centrifuged (5 min at 1,800 rcf, 4 °C) and washed once with 8 mL of Buffered Methanol-complex Medium (BMMY). After another centrifugation step (5 min at 1,800 rcf, 4 °C), the pellet was resuspended in 5 mL of BMMY medium and added to 15 mL of the same medium in 100 mL flasks, to obtain a 20 mL final volume. The cultures were then incubated at 30 °C and shaken at 220 rpm for 48 h. Every 24 h, methanol was added to a final concentration of 0.5 %, to promote expression of the heterologous protein under control of the methanol-inducible *AOX1* promoter. The same OD_600_ was harvested for each culture by centrifugation for 5 min at 1,800 rcf and stored at −80 °C until lipid extraction.

### *PpSD8D* subcellular localization in *P. patens via* particle bombardment

The *PpSD8D* coding sequence was amplified from cDNA and cloned into two pUC18-derived pEntry vectors (provided by Dr Ellen Hornung) containing a 35S promoter and an eYFP tag either N-or C-terminal to the multiple cloning site (primers in Supplemental Table S2). An ER marker plasmid, SP-mCerulean-KDEL, was obtained from Prof. Ralf Reski (Mueller & Reski, 2015). The bombardment followed the general protocol described in (Wegner *et al*., 2025). Four-week-old gametophores were used. Bombardment was performed by particle gun (PDS-1000/He; Bio-Rad, Hercules, USA) with 900 psi helium gas. Plates were placed four levels below the rupture disc and two levels below the macrocarrier. After bombardment, gametophores were cultivated for 48 h before imaging. For cell imaging, a Zeiss LSM 780 AxioObserver with LCI Plan-Neofluar 63x/ 1.3 mm Korr DIC M27 objective with a pinhole size of 1.00 AU was used. A 488 nm laser at 2.2 % intensity was used to excite eYFP, and detected between 506 to 552 nm. The 405 nm laser at 2 % intensity was used excite mCerulean, and signals were detected between 438-496 nm.

### Morphological phenotype analyses

Colonies were imaged using an Olympus SZX12 stereomicroscope with a Retiga R6 camera. Plates were ventilated under a sterile bench before imaging to reduce condensation. Gametophore colony area was measured, and together with colony perimeter used to calculate circularity according to the formula:

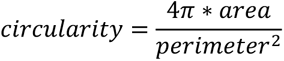

### Counting and measurement of phyllid cells using Cellpose

Single phyllids were dissected from two-month-old gametophores, destained in a 3:1 ethanol-acetic acid solution (v:v) until they were fully discolored, rinsed 3X with water, and finally stained in a 0.01 % (w/v) toluidine blue solution to visualize cell walls, and again rinsed 3X. Phyllids were imaged with an Olympus BX51 light microscope with a Retiga R6 camera. In the case of phyllids that exceeded the field of view of one image, multiple images were acquired and subsequently stitched in Adobe Illustrator.

For high-throughput analysis of cell number and geometry, an automated pipeline was established using Cellpose (Stringer *et al*., 2021). The pre-trained cyto3 model was refined by custom training using images of *P. patens* phyllids. The following parameters were applied based on iterative testing: flow threshold 0.6, and cell probability threshold −1.5. A custom Python-based pipeline was developed to integrate the Cellpose library, perform segmentation on all images to count cells, and measure, to scale in micrometers, the following parameters for each cell: area, centroid position, major axis length, and minor axis length.

### Phytohormone extraction

Phytohormone extractions were done with methyl-*tert*-butyl ether (MTBE) as described (Herrfurth & Feussner, 2020). 20 mg lyophilized material was ground using a Mixer Mill MM200 with steel beads for 30 s before being resuspended in 0.75 mL methanol. The suspensions were transferred to an 8 mL glass tube and mixed with 2.5 mL MTBE and internal standards (10 ng D_5_-jasmonic acid (JA), 30 ng D_5_-12-*oxo*-phytodienoic acid (OPDA), 10 ng D_5_-salicylic acid (SA)) before shaking for 1 h in a cold room at ∼ 6 °C. 0.6 mL water was added, then the samples were vortexed and left for 10 min at room temperature. After 10 min of centrifugation at 805 rcf, the upper phase of each sample was transferred to a clean glass tube and a second extraction was performed on the remaining lower phase, using 0.7 mL of methanol/water (3:2.5; v:v) and 1.3 mL of MTBE. The incubation and centrifugation steps were repeated and both phases without cell debris were added to the saved upper phase for evaporation under a nitrogen stream. The dry residues were then resuspended in 0.2 mL methanol and transferred to a 1.5 mL Eppendorf tube for evaporation. The extracts were finally dissolved in 20 µl B solvent (H_2_O:acetonitrile, 1:9 v:v, + 0.3 mM NH_4_HCO_2_, pH 3.5), vortexed and mixed with 80 µL A solvent (H_2_O with 0.3 mM NH_4_HCO_2_, pH 3.5). The tubes were then strongly vortexed and centrifuged for 5 min at 12,850 rcf. The supernatants were used for UPLC-nanoESI-MS/MS analysis. Before each incubation step, the extract was covered with argon gas.

### Phytohormone analysis

Phytohormones were separated on an ACQUITY UPLC system with an HSS T3 column (100 mm x 1 mm, 1.8 µm thickness), and ionized with a Triversa Nanomate nanoESI chip with 5 µm internal diameter nozzles; samples were measured on an AB Sciex 4000 QTRAP tandem mass spectrometer (Herrfurth & Feussner, 2020). Phytohormones were measured in negative ionization mode. Where appropriate internal standards were available (here, for salicylic acid and OPDA), quantifications were performed using an internal calibration curve based on the mass to charge (m/z) ratio detected vs. molar amount of standard injected. Where standard was not available, phytohormones were calculated as relative amounts adjusted to the dry weight. *tn*-OPDA was measured with a mass transition of 235/163, Δ^4^-*dn-*OPDA with a transition of 261/163, SA with a transition of 137/93, and OPDA with a transition of 291/165.

### Lipid extraction

The extraction described in (Herrfurth *et al*., 2021) was used for both *K. phaffii* and *P. patens* with minor modifications. In the case of *P. patens*, lipid extractions used 20 mg of lyophilized protonema and gametophores. Tissues were crushed with steel beads in a Mixer Mill MM200 before transferring to 8 mL glass tubes, and resuspending in 6 mL extraction buffer (propan-2-ol/hexane/water (60:26:14, v:v:v)). *K. phaffii* cells harvested according to OD_600_ were placed in 8 mL glass tubes containing 6 mL of extraction buffer and 0.5 mL glass beads and shaken for 30 min at 4 °C. From here, the methods are the same: the samples in extraction buffer were shaken at 60 °C for 30 min, with intermittent vortexing and sonication for 1 min every 10 min. The samples were then centrifuged for 20 min at 20 ⁰C at 805 rcf, the supernatants were transferred to clean glass tubes, and evaporated under a nitrogen stream. The extracts were resuspended in tetrahydrofuran/methanol/water (4:4:1, v:v:v), vortexed and sonicated for 1 min, before transferring to glass vials with an insert. Before every incubation and storage step the samples were covered with argon gas.

### Lipid Analysis

Sphingolipids were separated on an ACQUITY UPLC system with an HSS T3 column (100 mm x 1 mm, 1.8 µm thickness), and ionized with a Triversa Nanomate nanoESI chip with 5 µm internal diameter nozzles; samples were measured on an AB Sciex 6500 QTRAP tandem mass spectrometer (Herrfurth *et al*., 2021). Sphingolipids were measured in positive ionization mode, with [M+H]^+^ as the precursor ions for all lipid classes. The product ions were dehydrated LCBs for the free LCBs, ceramides, and GlcCers. The loss of the phosphoinositol-containing head group was used for the detection of GIPCs. MRM peak areas were corrected for loss of the ^13^C isotope. All signals were normalized to the amount of FAME detected per sample, as determined by gas chromatography with flame ionization detection (GC-FID) (Haslam *et al*., 2024).

## Results

### Pp6c16_5700V6.1 is an LCB Δ8-desaturase

A single ceramide backbone, 18:2;2/20:0;1, makes up approximately 95 % of the total GlcCer pool in *P. patens* (Gömann *et al*., 2021a; Resemann *et al*., 2021). The 18:2;2 LCB moiety is not detected in GIPCs, and is present in only trace amounts of free ceramides (Gömann *et al*., 2021a). In characterizing the genes most closely-associated with GlcCer metabolism in *P. patens* (Figure 1A), we built on the identification of *PpGCS* and *PpSD4D* carried out by (Gömann *et al*., 2021a). Further, a single *PpSD8D* gene candidate (Table 1) was identified by deduced amino acid sequence similarity to the two characterized *A. thaliana SD8D* paralogs, *AtSLD1* (70.57 % similarity to *PpSD8D*) and *AtSLD2* (78.5 % similarity to *PpSD8D*).

**Figure 1:**
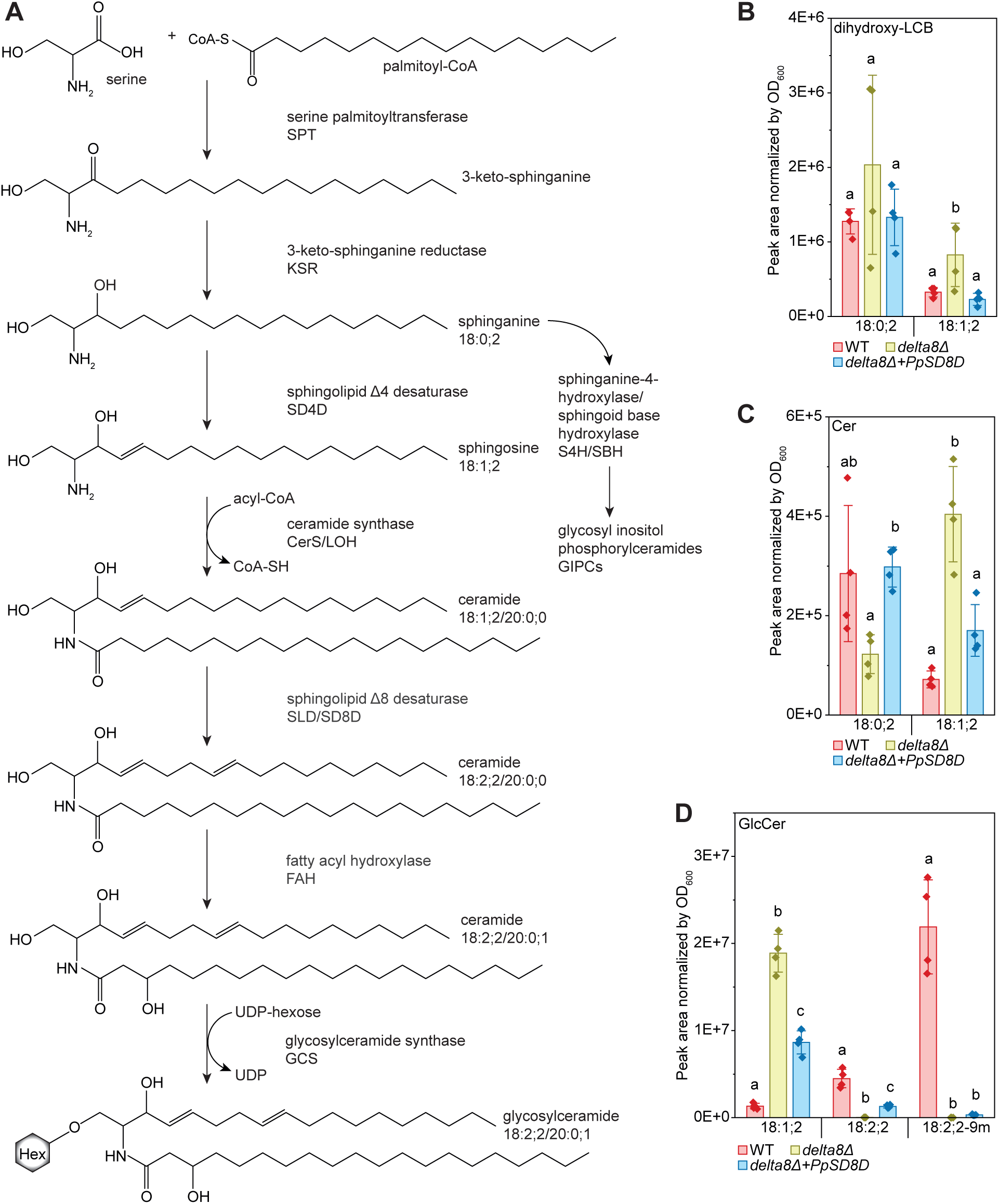
Synthesis of GlcCer in Embryophytes. (A) Proposed metabolic pathway. The reaction order is not absolutely known, in particular the sequence of SD4D and GCS activity; see discussion. (B-D) *Physcomitrium patens* SD8D activity demonstrated by heterologous expression and complementation of the *Δ8-desaturase* (*delta8Δ*) mutant of *Komagaetella phaffii*, sphingolipids were measured by UPLC-nanoESI-MS/MS. P. patens SD8D expression affects LCB (B), ceramide (C), and GlcCer (D) profiles, and is specifically evidenced by the partially restored accumulation of 18:2;2-containing GlcCer (D). Four replicates of each line (in the case of the complementation line, four independent transformed lines) were analyzed, data represent the mean ± SD. Statistical analysis was done using a one-way ANOVA with Tukey’s post-hoc test. Letters indicate significance at p < 0.05. Activities of *P. patens* SD4D and GCS were previously demonstrated (Goemann et al., 2021).

The predicted biochemical activity of *PpSD8D* was tested by complementation of the *K. phaffii* mutant *Δ8-desaturaseΔ* (*delta8Δ*) (Ternes *et al*., 2011)(Figure 1B-D). Similar to *P. patens*, *K. phaffii* preferentially incorporates 18:2;2 LCB, and Δ9-methylated 18:2;2-9m (fungi-specific) into their GlcCer (Ternes *et al*., 2011). While the *K. phaffii delta8Δ* mutant is indistinguishable from the wild type in its growth and general appearance, it completely lacks 18:2;2 and 18:2;2-9m sphingolipids, which is most obvious in GlcCers (Figure 1D). The mutant also accumulates sphingolipids containing an 18:1;2 LCB moiety compared to the wild type; this is observed in free LCBs (Figure 1B), free ceramides (Figure 1C), and GlcCers (Figure 1D). When *PpSD8D* was heterologously expressed, the amounts of the 18:1;2 LCB moiety in free LCBs and free ceramides were restored to wild-type levels (Figure 1B-C) and reduced in GlcCers (Figure 1D). Additionally, 18:2;2 LCB moieties were again detected in GlcCers, indicating that *PpSD8D* is a functional Δ8-desaturase. However, 18:2;2-containing GlcCer were less abundant than in wild type, and there was no recovery of 18:2;2-9m (Figure 1D), indicating *P. patens* SD8D activity in *K. phaffii* is inferior to the endogenous *Δ8-DESATURASE*, perhaps due to differences in gene product expression level or stability, stereochemistry of the double bond introduced, or compatibility with the sphingolipid-C9-methyltransferase. Localization of *Pp*SD8D to the ER, predicted based on other enzymes associated with ceramide assembly and modification, was confirmed by tagging the *PpSD8D* coding sequence at its N-terminus with eYFP, and transient expression of the construct in *P. patens* phyllid cells after particle bombardment (Supplemental Figure S1). Tagging *Pp*SD8D at its C-terminus produced very weak expression that we could not confidently interpret for localization.

### *sd4d gcs* double mutants have a similar sphingolipid profile to *gcs*, but accumulation of 18:2;2-containing free ceramides is suppressed

Having identified the *SD4D*, *SD8D*, and *GCS* genes in *P. patens*, we collected, generated, and characterized single and double mutants: *sd4d*, *sd8d*, *sd4d sd8d*, *gcs*, and *sd4d gcs*. Briefly, our previous characterization of protonema of *sd4d* single mutants uncovered a total absence of mono- and di-unsaturated LCB moieties in sphingolipids (apparent in both free ceramides and GlcCers), and a >99 % depletion in total GlcCers (Gömann *et al*., 2021a). This suggested that Δ4-desaturation is essential for both Δ8-desaturation and for GlcCer synthesis in *P. patens*. Characterization of the *gcs* single mutant revealed a very slightly more severe reduction in GlcCers, an accumulation of free ceramides containing 18:2;2 LCB moieties, and an increase in the amount of hydroxyceramides (OH-Cer, referencing the α-hydroxylation of the FA moiety) (Gömann *et al*., 2021a).

We generated *sd4d gcs* double mutants, *sd8d* mutants, and *sd4d sd8d* double mutants. As each genotype was isolated, we performed lipidomic analyses with three or more independent alleles of the new genotype of interest. Each of the individual analyses is presented in the supplement. For simplicity only one representative allele, from the final measurement in which all genotypes were available, is presented in the main Figure 2 for gametophores, and Figure 3 for protonema.

**Figure 2:**
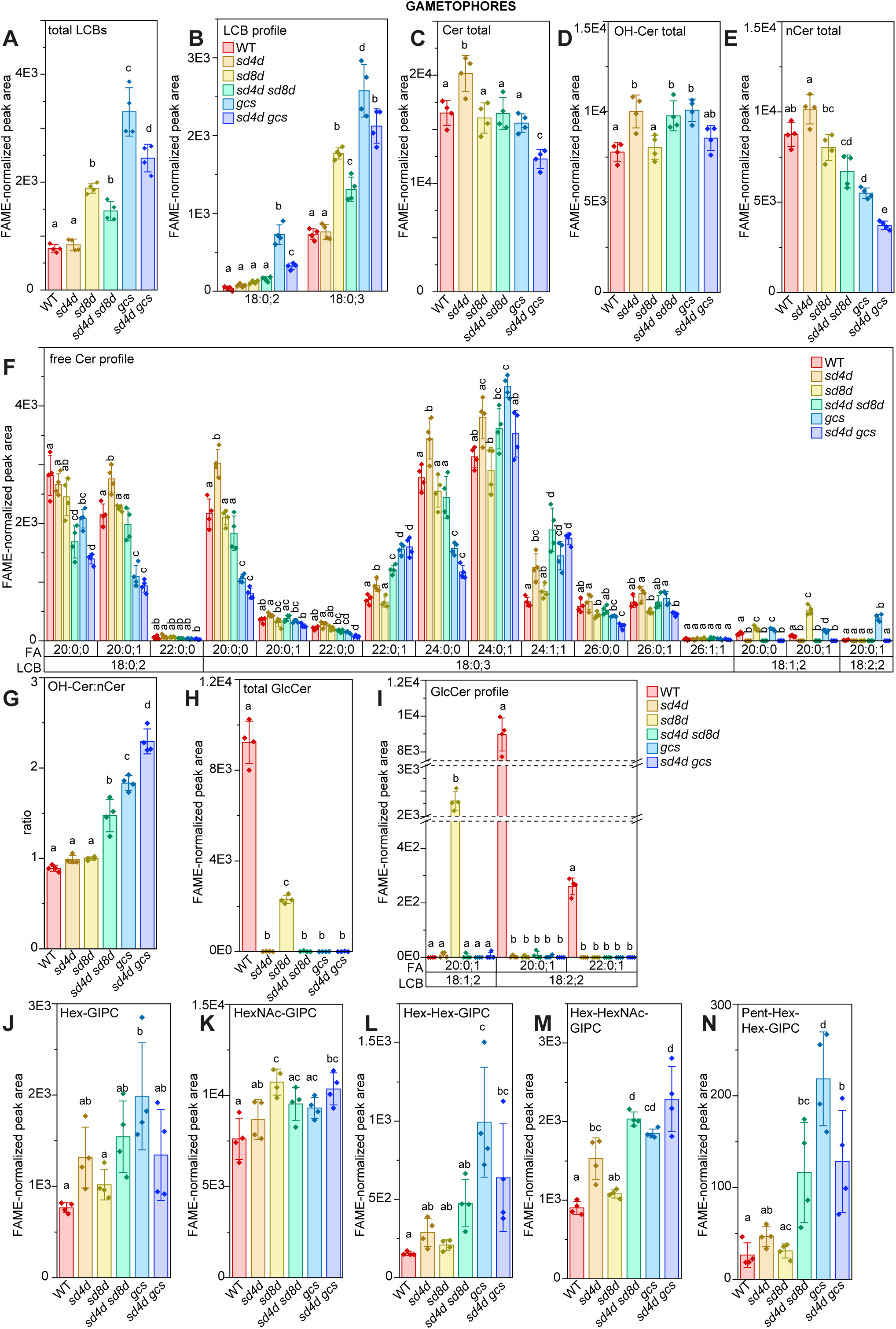
Sphingolipid profile of WT, *sd4d* (−1), *sd8d* (B1), *sd4d sd8d* (18), *gcs* (−3), and *gcs sd4d* (i11) gametophores. (A) Free LCB totals (B) Free LCB profile (C) Free ceramide totals (D) hydroxylated ceramide totals (E) non-hydroxylated ceramide totals (F) free ceramide profile (G) OH-Cer:nCer ratio (H) GlcCer total (I) GlcCer profile (J) Hex-GIPC total (K) HexNAc-GIPC total (L) Hex-Hex-GIPC total (M) Hex-HexNAc-GIPC total (N) Pent-Hex-Hex-GIPC total determined by UPLC-nanoESI-MS/MS. The peak areas are corrected for exclusion of the ^13^C isotope and normalized to the total FAMEs content of the sample. Data represent the mean ± SD of four replicates grown on separate plates. Statistical analysis was done using a one-way ANOVA with Tukey’s post-hoc test. Letters indicate significance at p < 0.05. Supported by Supplemental Table S3. WT: wild-type, *sd4d*: sphingolipid Δ4 desaturase, *sd8d*: sphingolipid Δ8 desaturase, *gcs*: glycosylceramide synthase, UPLC-nanoESI-MS/MS: ultra high-performance liquid chromatography with nanoelectrospray ionization and triple quadrupole mass spectrometry, FAME: fatty acid methyl ester, FA: fatty acid, LCB: long-chain base, GIPC: glycosyl inositol phosphorylceramide.

**Figure 3:**
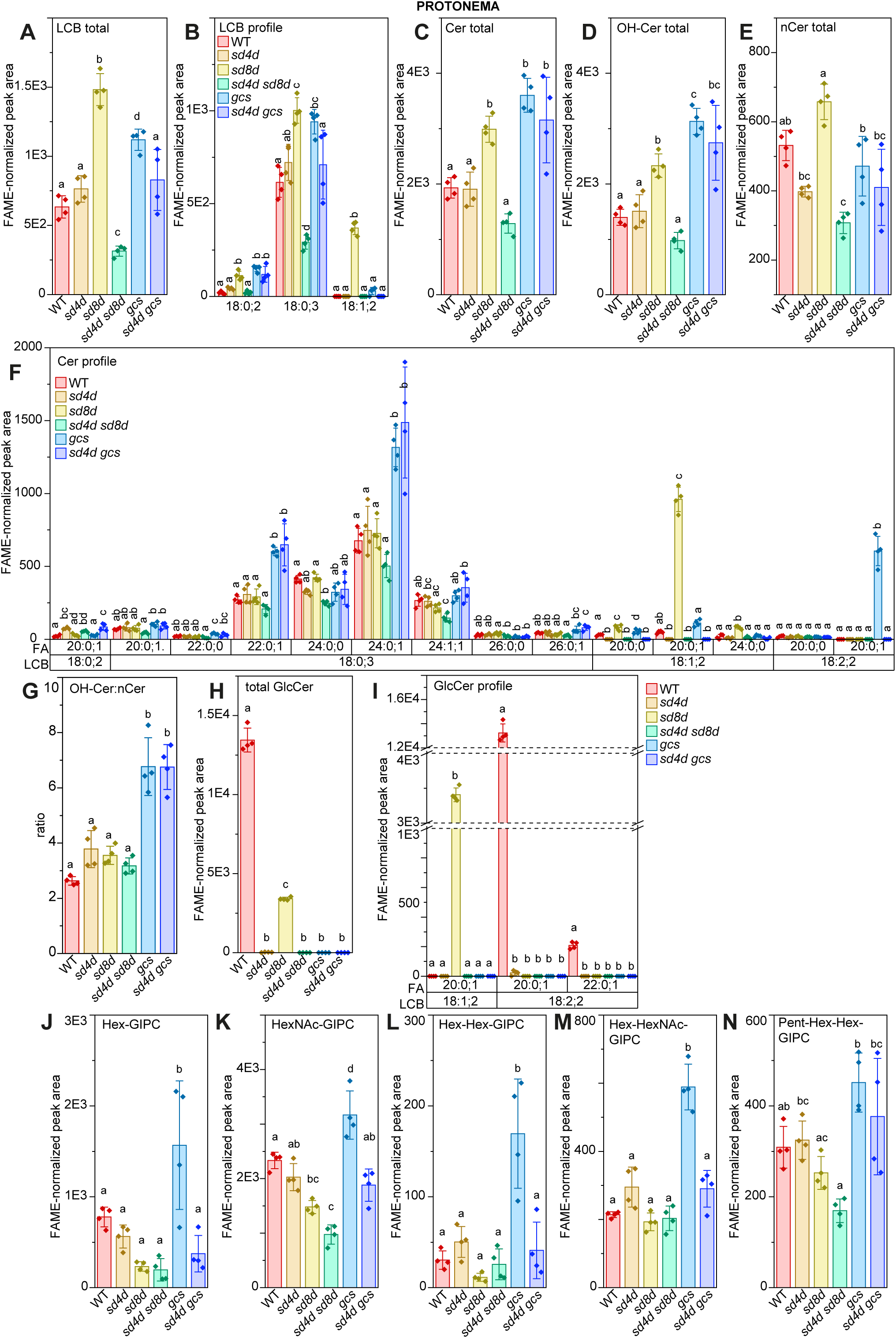
Sphingolipid profile of WT, *sd4d* (−1), *sd8d* (B1), *sd4d sd8d* (18), *gcs* (−3), and *gcs sd4d* (i11) protonema. (A) Free LCB totals (B) Free LCB profile (C) Free ceramide totals (D) hydroxylated ceramide totals (E) non-hydroxylated ceramide totals (F) free ceramide profile (G) OH-Cer:nCer ratio (H) GlcCer total (I) GlcCer profile (J) Hex-GIPC total (K) HexNAc-GIPC total (L) Hex-Hex-GIPC total (M) Hex-HexNAc-GIPC total (N) Pent-Hex-Hex-GIPC total, determined by UPLC-nanoESI-MS/MS.The peak areas are corrected for exclusion of the C13 isotope and normalized to the total FAMEs content of the sample. Data represent the mean ± SD of four replicates grown on separate plates. Statistical analysis was done using a one-way ANOVA with Tukey’s post-hoc test. Letters indicate significance at p < 0.05. Supported by Supplemental Table S4. WT: wild-type, *sd4d*: sphingolipid Δ4 desaturase, *sd8d*: sphingolipid Δ8 desaturase, *gcs*: glycosylceramide synthase, UPLC-nanoESI-MS/MS: ultra high-performance liquid chromatography with nanoelectrospray ionization and triple quadrupole mass spectrometry, FAME: fatty acid methyl ester, FA: fatty acid, LCB: long-chain base, GIPC: glycosyl inositol phosphorylceramide

In gametophores, *sd4d gcs* double mutants presented a similar chemotype to the single *gcs* mutant (Figure 2, Supplemental Figure S2). Both showed loss of GlcCers (Figure 2H-I, Supplemental Figure S2H-I). Critically, however, the loss of GlcCers in *gcs* singles, *sd4d gcs* doubles, and also *sd4d* singles was complete in our measurements (Figure 2, Figure 3, Supplemental Figures S2-S4). Therefore, the slightly more severe total GlcCer depletion previously reported (Gömann *et al*., 2021a) is not absolutely consistent, and is unlikely to be the cause for the difference in growth phenotypes of *sd4d* and *gcs*. As expected, however, the *sd4d gcs* double mutants differed from *gcs* in lacking free ceramides with monounsaturated and di-unsaturated LCB moieties (Figure 2F, Supplemental Figure S2F).

Additionally, both *gcs* and *sd4d gcs* had a 3-4-fold increase in total free LCBs (Figure 2A-B; Supplemental Figure S2A-B), and an approximately 2-fold increase in the OH-Cer to non-hydroxylated ceramides (nCer) ratio (OH-Cer:nCer) (Figure 2C-G). The persistent and even stronger increase in the OH-Cer:nCer ratio in *sd4d gcs* compared to *gcs* may be surprising, given that the simplest explanation for the increased ratio in *gcs* single mutants would be the accumulation of 18:2;2;/20:0;1 OH-Cer precursors of GlcCer. The increased ratio in *sd4d gcs* indicates that there are broader shifts in the free ceramide profile in *gcs* that are not directly related to SD4D catalytic activity. Finally, both *gcs* and *sd4d gcs* accumulated more GIPCs, especially the more highly-glycosylated B- and C-series GIPCs, which were increased 2-10-fold in these two genotypes. (Figure 2J-N, Supplemental Figure S2J-N).

Analysis of the same mutant genotypes grown as protonema was also performed, for direct comparison to our previous results (Gömann *et al*., 2021a) (Figure 3A-N). This revealed largely similar trends to the gametophore measurement described above, except that the GIPC accumulation observed in *gcs* single mutants was not retained in the *sd4d gcs* double mutants (Figure 3J-N). In both gametophores and protonema, however, the most anticipated chemotypes for *sd4d gcs* were conserved; the loss of GlcCers was consistent with that observed in *gcs* and *sd4d* single mutants, while the accumulation of free ceramides containing 18:1;2; and 18:2;2 LCB moieties characteristic of the *gcs* single mutant was suppressed.

### The *sd8d* mutant chemotype supports the hypothesis that Δ8-desaturation occurs on a Δ4-desaturated substrate

*sd8d* single mutants were generated and analyzed with the expectation that they would affect the synthesis of GlcCers downstream of SD4D activity. (Figure 2A-N; Supplemental Figure S3A-N). Overall the free ceramide, OH-Cer, and nCer levels, as well as the ratio of OH-Cer:Cer were not substantially affected in the *sd8d* mutant (Figure 2C-G; Supplemental Figure S3A-G). The free ceramide profile of *sd8d* only meaningfully and significantly differed from the wild type in that species containing 18:1;2 LCB moieties accumulated (Figure 2F; Supplemental Figure S3F), fitting with the expectation that SD8D accepts and desaturates the monounsaturated products of SD4D activity prior to their integration into GlcCers. Unexpectedly, *sd8d* mutants did seem to accumulate free LCBs (Figure 2A-B), however, this was not reproducible across measurements and independent alleles (Supplemental Figure S3A-B).

Most prominently, the total GlcCer amount in *sd8d* mutants was reduced by approximately two thirds compared to the wild type, and consisted entirely of 18:1;2/20:0;1 (Figure 2H-I; Supplemental Figure S3H-I). While substantial and significant, this depletion is far less dramatic than that of the *sd4d* single mutants, suggesting that for GlcCer production in *P. patens*, Δ8-desaturation is either less critical than Δ4-desaturation, or ceramides containing monounsaturated LCBs are better substrates for glycosylation by GCS than ceramides containing fully saturated LCBs. Finally, the GIPC contents of the *sd8d* mutants were not substantially or consistently different from the wild type (Figure 2J-N; Supplemental Figure S3J-N).

Compared to gametophores, analysis of sphingolipids in *sd8d* protonema (Figure 3A-N) was consistent in that the loss of Δ8-desaturation produced an approximately two thirds depletion in GlcCers (Figure 3H), and accumulation of 18:1;2 LCB moieties in free ceramide form and in GlcCer. Yet, there were two substantial differences between the gametophore and protonemal chemotypes. First, free LCB measurement in protonema revealed accumulation of 18:1;2 species (Figure 3B). This was a surprising finding given that normally only saturated free LCBs can be detected in *P. patens*, and meaningful given that the reaction order for LCB moiety desaturations and ceramide synthesis are ambiguous (Haslam & Feussner, 2022). The appearance of 18:1;2 in the free LCB pool suggests that either SD4D accepts free LCB substrates, or that in the case the substrate of SD4D is a free ceramide, its 18:1;2 product undergoes degradation to free LCBs when SD8D activity is lacking. Second, the accumulation of 18:1;2 LCB-containing free ceramides was far more pronounced in the profile of protonema, with 18:1;2/20:0;1 becoming the single most abundant free ceramide, representing approximately one third of the total free ceramide pool in *sd8d* (Figure 3F). In contrast, in gametophores, 18:1;2 LCB-containing free ceramides remained minor species representing approximately 6 % of the total free ceramides in *sd8d* (Figure 2F). The magnitude of the 18:1;2 LCB-containing free ceramide increase in protonema resulted in an overall 50 % increase in total free ceramide amount (Figure 3C). Yet, unlike the free ceramide shifts observed in *gcs* and *sd4d gcs*, this did not have a significant effect on the overall OH-Cer:nCer ratio (Figure 3G).

### *sd4d sd8d* double mutants lack desaturated LCB moieties in all sphingolipids, and display other general shifts in free LCB and free ceramide profiles

Analysis of *sd4d sd8d* double mutants was anticipated to reveal a similar chemotype to *sd4d* singles, given the observed dependence of SD8D on SD4D activity. This was generally true, with minor exceptions (Figure 2, Figure 3, Supplemental Figure S4). First, relative to the wild type, the free LCB levels of *sd4d sd8d* showed an approximately 50 % accumulation in gametophores (Figure 2A, Supplemental Figure S4A), and approximately 50 % depletion in protonema (Figure 3A), whereas free LCB levels were not significantly different from the wild type in *sd4d*. Second, in gametophores, there was a marked depletion in nCer that produced an increase in the overall OH-Cer:nCer ratio, which was not observed in *sd4d* (Figure 2E, 2G, Figure 3E, 3G, Supplemental Figure S4E, S4G). Otherwise, the *sd4d* and *sd4d sd8d* chemotypes were indeed similar.

### Developmental phenotypes of *gcs* are not suppressed when ceramide substrate accumulation is blocked in the *sd4d gcs* double mutant

Lipidomic analysis determined that *sd4d gcs* double mutants present a similar chemotype to *gcs* singles in gametophores, with the exception that the accumulation of free ceramides containing 18:1;2 and 18:2;2 LCB moieties was suppressed. Therefore, the growth phenotype of the double mutants should indicate whether the severe growth phenotype of *gcs* single mutants is due to accumulation of these specific and unusual free ceramides. Across all developmental stages observed, *sd4d gcs* doubles were as stunted as the *gcs* single mutants (Figure 4). Gametophore colonies of *sd4d gcs* appeared as impaired as *gcs* (Figure 4A), and this could be quantitatively confirmed by colony area measurements finding that *gcs* and *sd4d gcs* double mutants were both reduced to one sixth that of the wild type (Figure 4B), and colony circularity increased by 2-2.5-fold (Figure 4C).

**Figure 4:**
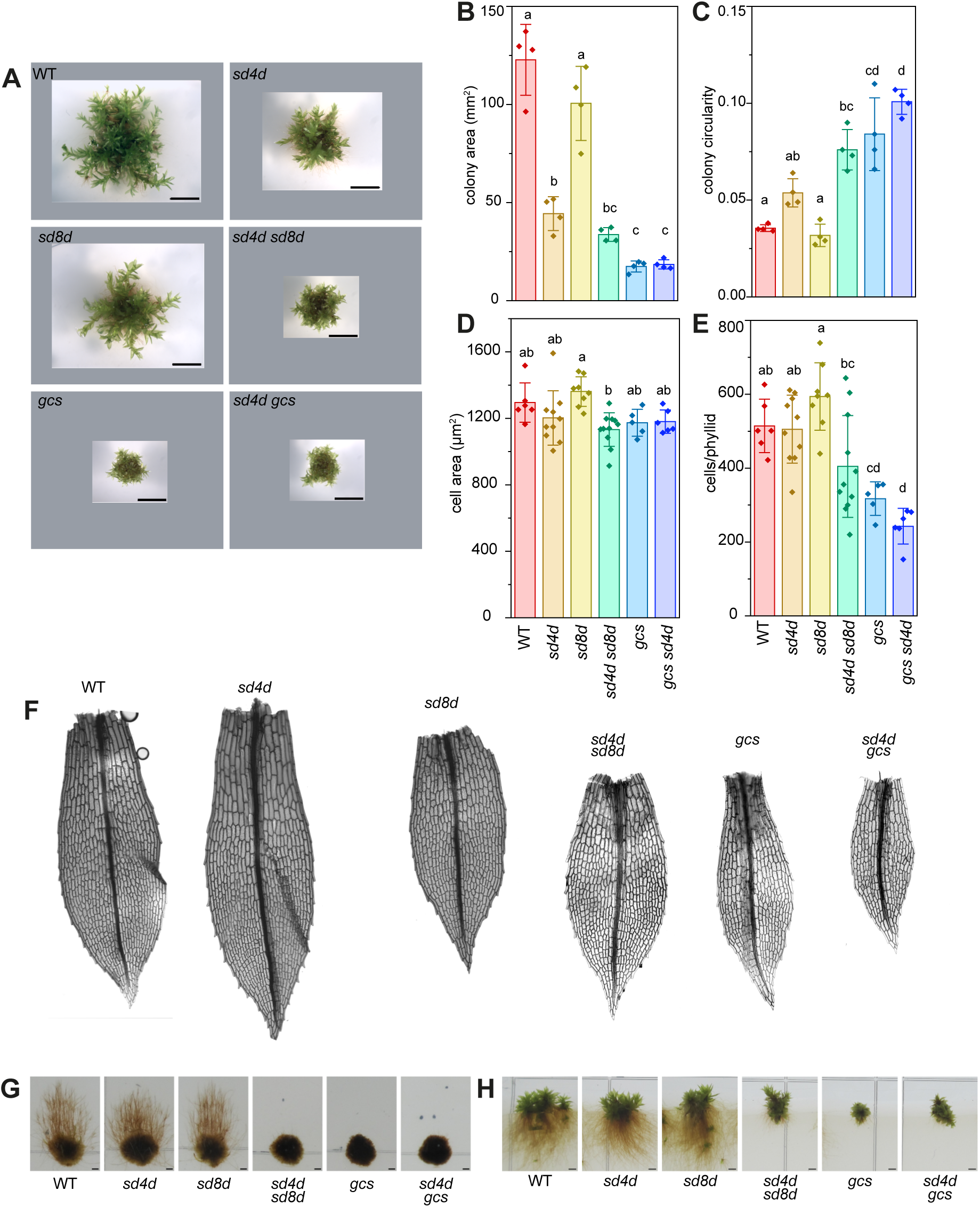
Morphological phenotypes of the mutants. (A) Images of 48-day old gametophore colonies. (B) Colony area. (C) Colony circularity. For B and C, bars represent the averages of four colonies, error bars represent standard deviation. (D) Average phyllid cell area. (E) Average number of cells per phyllid. For D and E, data points represent average values calculated from five to eleven individual phyllids, bars are the averaged values of all phyllids for each genotype, and error bars represent standard deviation. For all plots, letters indicate significance at p < 0.05 using a one-way ANOVA with Tukey’s post-hoc test. (G) Skotonemal cultures, i.e. protonema grown vertically in the dark on sucrose-supplemented media. (H) Rhizoid cultures. For G and H, all genotypes were grown on the same plate with identical treatments and conditions, but images were cropped for consistent ordering in data presentation. All scale bars represent 4 mm. WT: wild type, *sd4d*: sphingolipid Δ4 desaturase, *sd8d*: sphingolipid Δ8 desaturase, *gcs*: glycosylceramide synthase, KD: knock-down, KO: knock-out.

Analysis of single phyllids (Figure 4D-F) revealed no substantial and consistent differences in the average cell area of the genotypes measured (Figure 4D). In contrast, the number of cells per phyllid showed significant reductions in both the *gcs* and *sd4d gcs* mutants (Figure 4E). The magnitude in the reductions in cell number were not proportional to the reduction in colony area, suggesting that other factors, such as the number of gametophores developed, gametophore cauloid elongation, and phyllid organogenesis, could impact the overall growth phenotype.

Phenotypes were also observed in other growth stages. Skotonemal growth, which we previously observed was non-existent in *gcs*, was also absent in *sd4d gcs* (Figure 4G). There was a similar, obvious deficiency in rhizoid growth in the *gcs* and *sd4d gcs* mutants (Figure 4H). Altogether, these phenotypes consistently indicate that the suppressed accumulation of 18:1;2 and 18:2;2 free ceramides in *sd4d gcs* does not rescue the growth phenotype of *gcs*. Other elements of the *gcs* mutant chemotype, especially other shifts in the free ceramide profile, the OH-Cer:nCer ratio, or changes to LCB accumulation, must be contributing to *gcs* developmental phenotypes.

### *sd4d sd8d* development is more severe than *sd4d* single mutants, despite their similar chemotypes

We also observed and quantified the growth phenotypes of the *sd8d* and *sd4d sd8d* mutants. The *sd8d* single mutants were, in all growth stages observed and measured, indistinguishable and not significantly different from the wild type. In contrast, the *sd4d sd8d* double mutants appeared more severe than the wild type, and importantly, more severe than *sd4d* singles. We first observed this visually in the gametophore stage (Figure 4A), although the colony and cell measurements showed no significant difference (Figure 4B-F). However, skotonemal and rhizoid growth in *sd4d sd8d* doubles was completely abolished, as in the *gcs* and *sd4d gcs* mutants (Figure 4G-H). Given (1) the similarity of the *sd4d* and *sd4d sd8d* mutant sphingolipid chemotypes, and (2) our understanding that *sd4d* should be epistatic to *sd8d* based on the phenotypes of the single mutants, we had fully expected that *sd4d sd8d* should resemble *sd4d* single mutants. This difference in *sd4d sd8d* indicates that *sd4d* is not fully epistatic to *sd8d*; it could be that additional shifts in the free LCB and free Cer profiles, or even additional functions of the *SD8D* gene product, contribute to these growth phenotypes.

### Cold stress exacerbates the *gcs* mutant phenotype, and results in up-regulation of transcripts associated with oxylipin metabolism

We noticed that the growth phenotypes of the *gcs* mutant were exacerbated in the cold conditions we use to maintain our backup strain collection (short days, 17 ⁰C, and low light intensity) (Figure 5A). We also observed the appearance of small, brown patches on *gcs* phyllids (white arrows in Figure 5A). These brown patches are reminiscent of lesion mimic phenotypes often observed in sphingolipid-deficient mutants of *A. thaliana* (reviewed in (Berkey *et al*., 2012)). Such lesion-mimic phenotypes are associated with hyperaccumulation of the phytohormone salicylic acid (SA)(König *et al*., 2022). SA is a key regulator of plant defense, especially against biotrophic pathogens.

**Figure 5:**
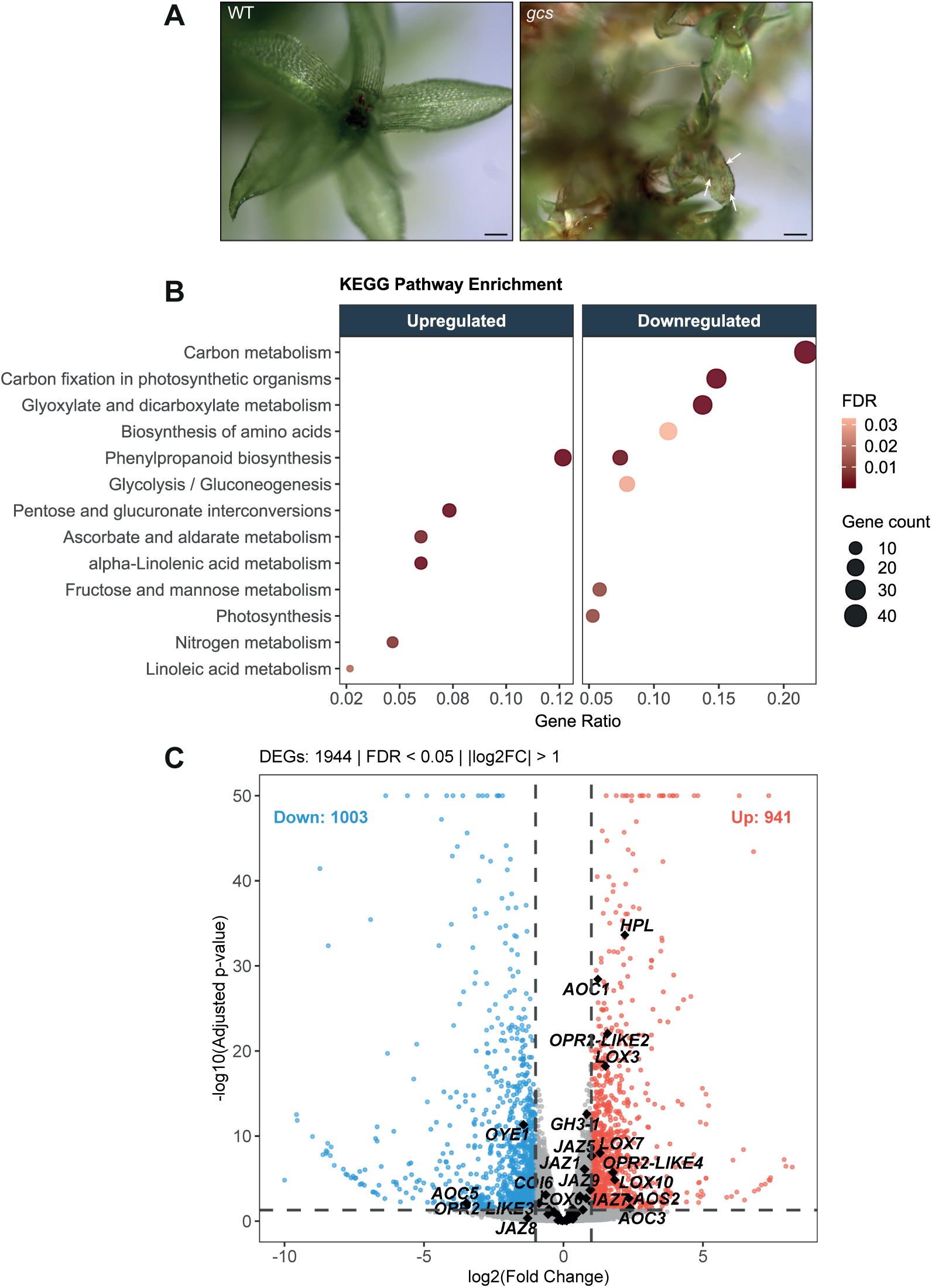
Cold stress exacerbates the phenotype of *gcs*, resulting in transcriptional differences between wild type and *gcs*. (A) Under cold stress, the *gcs* mutant develops brown patches (white arrows) on its phyllids. Scale bars are 2 mm. (B) Transcripts associated with different KEGG pathways are differentially expressed in the *gcs* transcriptome compared to the wild type, with carbon metabolism being strongly down-regulated, and phenylpropanoid metabolism, alpha-linolenic acid metabolism, and linoleic acid metabolism strongly up-regulated. (C) Multiple transcripts involved in oxylipin (i.e α-linolenic acid and linoleic acid) metabolism (bold) are up-regulated in *gcs*. *gcs*: glycosylceramide synthase, KEGG: Kyoto encyclopedia of genes and genomes, FDR: false discovery rate, DEG: differentially expressed gene, *AOC*: *ALLENE OXIDE CYCLASE*, *AOS*: *ALLENE OXIDE SYNTHASE*, *COI*: *CORONATINE INSENSITIVE*, *GH3*: *GRETCHEN HAGEN3*, *HPL*: *HYDROPEROXIDE LYASE*, *JAZ*: *JASMONATE ZIM DOMAIN*, *LOX*: *LIPOXYGENASE*, *OPR*: *OXO-PHYTODIENOIC ACID REDUCTASE*.

To explore the relationship between sphingolipid metabolism, phytohormone signaling, and the phenotypes of our *P. patens* mutants, we performed RNAseq on triplicate samples of the wild type and *gcs* mutant, grown under cold stress conditions (complete data: Supplemental Table S7, filtered and analyzed data in Supplemental Table S8). Of 20,979 detected transcripts, there were 941 up-regulated and 1003 down-regulated transcripts in the mutant, using a threshold of a log_2_ fold-change >1, and −log_10_(p value)<0.05 to identify differentially-expressed genes (DEGs). *GCS*, *SD4D*, and *SD8D* were not differentially expressed (Supplemental Table S9). Automatically-assigned KEGG pathway terms revealed an enrichment of transcripts associated with phenylpropanoid biosynthesis, α-linolenic acid (18:3n-3) and linoleic acid (18:2n-6) metabolism, pentose and glucuronate interconversions, and nitrogen metabolism (Figure 5B). The enrichment of transcripts associated with phenylpropanoid metabolism is consistent with brown pigmentation observed in cold-stressed *gcs*, and the enrichment of transcripts associated with α-linolenic acid and linoleic acid metabolism suggests that oxylipin synthesis could be differentially regulated in cold-stressed *gcs*. A targeted search for genes associated with oxylipin metabolism revealed strong and significant up-regulation of many of these (Figure 5C, Supplemental Table S9), supporting the notion that oxylipin signaling could be activated in cold-stressed *gcs*.

### Oxylipins accumulate in *gcs* mutants, and accumulation is exacerbated by cold treatment

We measured stress-associated phytohormones including SA and the cyclopentenones OPDA, Δ^4^-dn-OPDA, and tn-OPDA in the wild type and *gcs* grown in cold and control conditions (Figure 6A-D). We observed no significant difference in SA levels (Figure 6A), or OPDA levels (Figure 6B). The *cis*- and *iso*-isomers of OPDA could not be clearly and separately integrated, therefore the measurements combine both isomers. In contrast, Δ^4^-dn-OPDA (Figure 6C), and tn-OPDA (Figure 6D) showed clear enrichment in *gcs* compared to wild type, and this enrichment was strongly enhanced upon cold stress.

**Figure 6:**
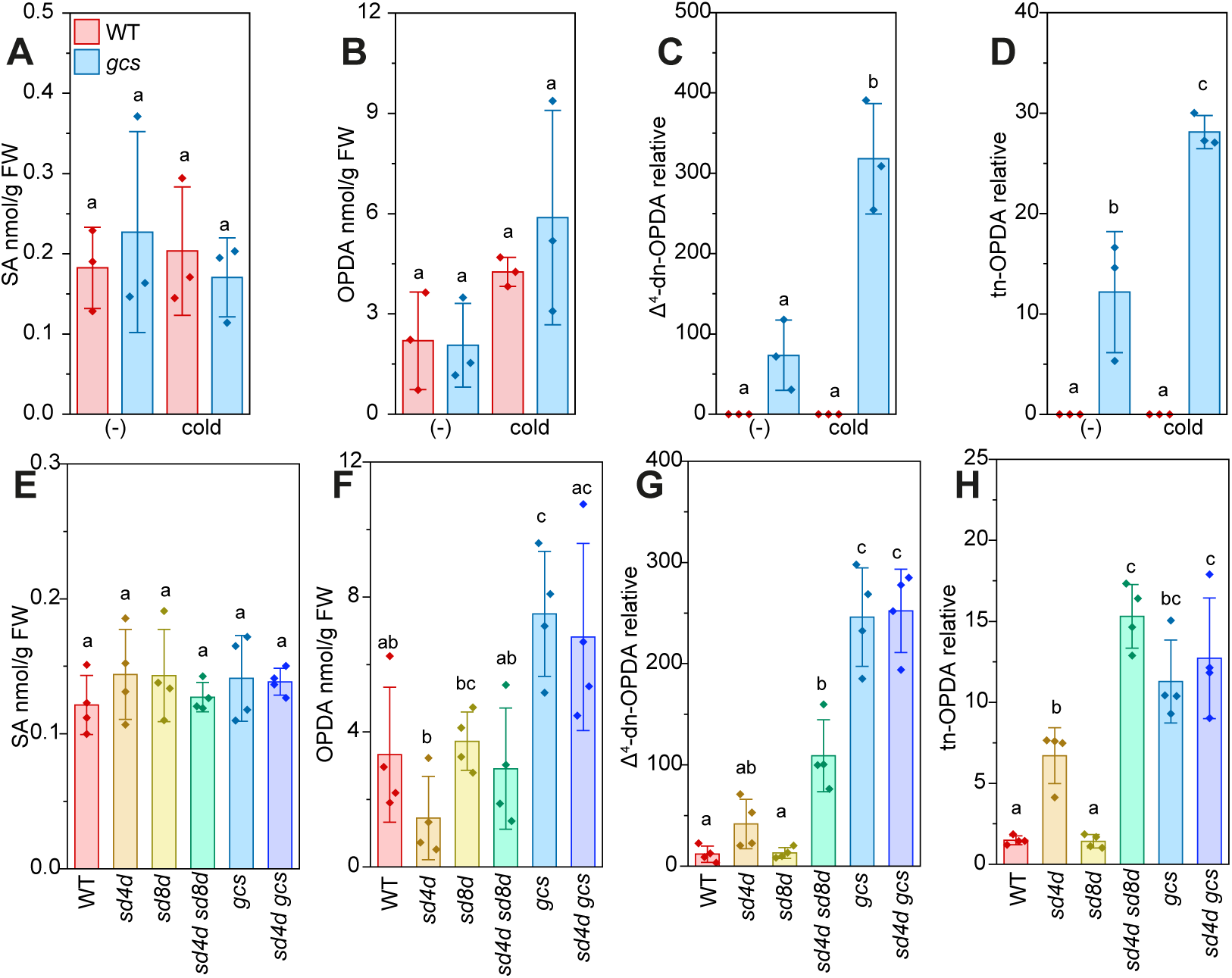
Cold stress and manipulation of GlcCer synthesis affects oxylipin metabolism. (A-D) Phytohormone measurements for WT and *gcs* (−3) grown under standard conditions (negative control, -), and under autumnal conditions for 6 weeks, that is, 17 °C and short days (8h light/16h dark). Data represent the mean ± SD of three replicates grown on separate plates. (E-G) Phytohormone measurements of WT, *sd4d* (−1), *sd8d* (B1), *sd4d sd8d* (18), *gcs* (−3), and *gcs sd4d* (i11) determined by UPLC-nanoESI-MS/MS. OPDA and SA are quantified in absolute values using a calibration curve, whereas Δ^4^-dn-OPDA and tn-OPDA are presented in relative amounts, in the absence of a comparable standard. ‘OPDA‘ represents both the *cis*- and *iso*- isomers, as both peaks could be detected but not clearly and separately integrated. Statistical analysis was done using a one-way ANOVA with Tukey’s post-hoc test. Letters indicate significance at p < 0.05. WT: wild type, *sd4d*: sphingolipid Δ4-desaturase, *sd8d*: sphingolipid Δ8-desaturase, *gcs*: glycosylceramide synthase, GlcCer: glycosylceramide, SA: salicylic acid, OPDA: 12-*oxo*-phytodienoic acid, FW: fresh weight.

We then measured cyclopentenones in all *gcs*, *sd4d*, and *sd8d* mutants to determine whether these accumulation patterns are consistent among genotypes similarly affected in GlcCer metabolism. SA was not significantly different in any genotype. Some accumulation of OPDA could be observed in *gcs* and *sd4d gcs*, but this was only significant for *gcs* (Figure 6E). Yet, for Δ^4^-dn-OPDA and tn-OPDA, the two cyclopentenones most strongly up-regulated in *gcs* in the cold, we could again observe a clear and significant increase in *sd4d sd8d*, *gcs*, and *sd4d gcs* (Figure 6F, 6G). These trends suggest that cyclopentenone synthesis is affected when GlcCer synthesis is disrupted. Further, the intensity of Δ^4^-dn-OPDA and tn-OPDA accumulation across genotypes and conditions correlates with the intensity of visually obvious signs of stress.

## Discussion

### Disrupted free Cer homeostasis is a likely cause of severe growth phenotypes in *gcs* and *sd4d gcs*

At the outset of this work, we hypothesized that the different developmental phenotypes of *sd4d* and *gcs* were either due to the previously observed more severe GlcCer depletion in *gcs*, or the accumulation of 18:2;2/20:0;1 free OH-Cer precursors of GlcCer in *gcs*. In the multiple lipidomic measurements we carried out here however, there was no significant difference in the levels of total GlcCer between *sd4d*, *gcs*, and *sd4d gcs*, demonstrating that the basis for our first hypothesis was not absolutely reproducible. This is not entirely surprising given the small amounts of GlcCer initially detected in these mutants, which are near the limit of detections for our methods. Additional factors that could have affected our lipidomic measurements include slightly different handling, MRM signal normalization methods to FAMEs instead of dry weight, and a change of light source from incandescent to LED. The *sd4d gcs* double mutants showed suppression of 18:2;2/20:0;1 free ceramide accumulation, as could be expected with the introduction of *sd4d*. Yet, because the growth phenotype of *sd4d gcs* was as severe as *gcs* alone, altogether these results refute both of our initial hypotheses.

Beyond these chemotypes directly affected by the metabolic steps mutagenized, we observed other significant changes in our sphingolipid measurements, most notably in the free ceramide profile. Interpreting these other shifts provides new hypotheses to explain the observed differences in growth phenotypes.

One of the most consistent differences between the sphingolipid profiles of *gcs* and *sd4d gcs* compared to *sd4d* was the OH-Cer:nCer ratio. From this, we speculate that the disruption of free ceramide homeostasis is complex in *gcs* mutants: not only do OH-Cer with 18:2;2 and 18:1;2 LCB moieties accumulate, but some OH-Cer containing 18:0;2 and 18:0;3-LCBs also accumulate, and there is a clear decrease in nCer in these mutants. This last trend persists when the accumulation of 18:2;2 and 18:1;2 LCBs is blocked by the introduction of *sd4d* into *gcs*. The significance of an increased OH-Cer:nCer ratio is not inherently obvious, as the accumulation of both have been associated with different physiological responses (Magnin-Robert *et al*., 2015; Zienkiewicz *et al*., 2020). The chemotype nevertheless indicates a broad disruption of ceramide metabolism.

Another less-obvious change is that many 18:0;3-containing free ceramides accumulate in *sd4d* gametophores(Figure 2F, Supplemental Figures S2, S3, S4), likely because the position of the third hydroxylation is the same as the Δ4-desaturation. This mutual exclusivity implies that loss of SD4D will shift the balance of ceramides incorporating 18:0;3 vs. 18:1;2 LCBs, because SPHINGANINE-4-HYDROXYLASE (S4H) and SD4D effectively compete for substrate. We grouped and summarized the free ceramides according to their LCB moiety (Supplemental Figure S6); while the total free ceramide amount varies substantially among genotypes, we observed a strong accumulation of free ceramides with 18:0;3 uniquely in *sd4d*. The amount of free ceramides with 18:0;3 is either unchanged or depleted relative to WT in both *gcs* and *sd4d gcs*. It is puzzling that the amount of 18:0;3 remains so low in *sd4d gcs*, however, the relative amount of 18:0;3 as a fraction of the total increases. Recent work producing a cryo-electron microscopy (cryo-EM) structure of serine palmitoyl transferase (SPT) (Liu *et al*., 2023; Cahoon *et al*., 2025) indicates that ceramides containing 18:0;3 LCBs are efficient negative regulators of SPT via the OROSOMUCOID-LIKE (ORM) component of the SPT complex. 18:0;2-containing ceramides have no inhibitory effect on SPT, and 18:1;2-containing ceramides have 1/10^th^ the effect on SPT as those with 18:0;3. We suggest that *sd4d* is unique among GlcCer-affected mutants, in that it restricts the severity of its own chemotype due to the negative feedback regulation of sphingolipid synthesis at SPT, promoted by the increased synthesis of 18:0;3-containing free ceramides when *SD4D* is knocked out. The accumulation of ceramides with 18:0;3 in *sd4d*, instead of accumulation of free ceramides with 18:1;2 (or 18:2;2) in *gcs*, could be one reason why *sd4d* accumulates less GIPCs—a possible sink for excess total ceramides. Enhanced negative feedback of total sphingolipid production may also be a reason why the *sd4d* phenotype is so much less severe than that of *gcs*. We expect this is one factor, in addition to GlcCer depletion, the OH-Cer:nCer ratio, and LCB accumulation, that affects the phenotypic spectrum of these mutants.

### The balance between GlcCers and GIPCs in different sphingolipid mutants is likely a consequence of disrupted free ceramide homeostasis, not compensation between the two complex sphingolipid classes

The two forms of complex sphingolipids present in plants, GlcCers and GIPCs, are defined by their distinct headgroups. Additionally, particular ceramide backbones can be associated, often strongly, with either GlcCers or GIPCs. This suggests that ceramide modifications contribute to partitioning within sphingolipid metabolism. Yet, disrupting LCB hydroxylation, a modification broadly associated with GIPC synthesis across analyzed plant species, has no negative effect on GIPC accumulation. In fact, *s4h/sbh* mutants of both *A. thaliana* (Chen *et al*., 2008) and *P. patens* (Gömann *et al*., 2021b; Steinberger *et al*., 2021) exhibit accumulation of GIPCs. This increase has been explained by the 18:0;3 direct product of S4H/SBH regulating the catalytic activity of SPT as described above, whereas the ceramides containing 18:0;2 that accumulate in *s4h/sbh* are ineffective (Steinberger *et al*., 2021; Cahoon *et al*., 2025). Downstream, the absence of 18:0;3 apparently does not interfere with the synthesis of GIPCs, as 18:0;2 fully replaces it—in both *A. thaliana* and in *P. patens*.

In contrast, desaturated LCBs are typically incorporated into GlcCers, and interfering with LCB desaturation limits flux into GlcCers in both *A. thaliana* (Δ8-desaturation) (Chen *et al*., 2012)and *P. patens* (both Δ4- and Δ8-desaturations) (Gömann *et al*., 2021a). Therefore, fundamentally different mechanisms regulate the synthesis of GlcCers vs. GIPCs—dependent and independent of LCB modifications, respectively.

Notably, in our sphingolipid measurements of gametophores of GlcCer-affected mutants, we observed an accumulation of GIPCs in the genotypes showing the strongest developmental phenotypes for this growth stage: *gcs*, *sd4d gcs*, and *sd4d sd8d*. While this superficially resembles a compensatory effect, this explanation conflicts with our understanding of the spatial distribution of GlcCers throughout the extra-plastidial membranes and enrichment in the tonoplast (Haschke *et al*., 1990; Andersson *et al*., 2005; Carmona-Salazar *et al*., 2021; Guo *et al*., 2022), in contrast to GIPCs in the plasma membrane (Bahammou *et al*., 2024). Compensatory effects of GlcCers and GIPCs are also questionable given the distinct physicochemical properties of these lipids in terms of the size and electrostatic charge of their headgroups, and their preferential incorporation of substantially different chain lengths of fatty acyl groups. We postulate that rather than compensation, the accumulation of GIPCs in *gcs*, *sd4d gcs*, and *sd4d sd8d* is a consequence of the dysregulation of SPT caused by the broad shifts in the free ceramide profile. In this case, GIPC synthesis could serve as a sink to remove excess free ceramides from the cell.

### Δ8 desaturation is less critical than **Δ**4 for ceramide flux into GlcCer in *P. patens*

We previously concluded that *Pp*SD8D is dependent on *Pp*SD4D for activity, based on the absence of any LCB desaturations in the *Ppsd4d* single mutant. Characterization of *Ppsd8d* single mutants supported this notion, as they presented strong accumulation of monounsaturated LCBs (presumably Δ4). Further, the presence of approximately one third of the normal GlcCer levels in *Ppsd8d* revealed that its activity contributes, but is not essential, for ceramide partitioning into GlcCer.

The remaining GlcCer here was entirely composed of 18:1;2/20:0;1 ceramide moieties, and this ceramide in free form also accumulated substantially: Especially in protonema, 18:1;2/20:0;1 became the single most abundant free ceramide, taking up approximately one third of the total pool. Despite these drastic modifications to both free ceramides and GlcCer, the *sd8d* single mutants remained in all measurable respects wild type-like. This presents an interesting similarity to findings in *A. thaliana*. Mutagenesis of the two *A. thaliana* genes encoding Δ8-desaturases (*AtSLD1* and *AtSLD2*) produced lines completely lacking the Δ8 desaturation, which is normally present as an 18:1;2 LCB moiety preferentially incorporated into GlcCer in *A. thaliana*. The double mutants also presented an approximately 50 % depletion in total GlcCer; despite this substantial chemotype, they only presented growth defects under cold stress (Chen *et al*., 2012). This resilience in both *P. patens* and *A. thaliana* to 30-50 % depletion in this complex sphingolipid class underlines the fact that while some elements of sphingolipid homeostasis are critical for normal development and physiology, others are either quite flexible, or non-essential under controlled laboratory conditions.

### Free ceramide homeostasis is disrupted in *sd4d sd8d*, despite SD4D appearing epistatic to SD8D in its direct activity

The isolation of *sd4d sd8d* double mutants was expected to produce an identical phenotype to *sd4d*, given the total absence of LCB desaturation in *sd4d*. While the double mutants did still present fully saturated LCBs, they differed from either single mutant in that they had an increased OH-Cer:nCer ratio, similar to *gcs* mutants. They also accumulated less free ceramides with 18:0;3 moieties compared to *sd4d* single mutants (Supplemental Figure S6). Both of these results suggest that although the direct catalytic activity of SD8D is dependent on SD4D, free ceramide homeostasis is somehow further disrupted when both *SD8D* and *SD4D* are knocked out. These could be related to regulatory, non-enzymatic functions, or perhaps an unknown secondary enzymatic activity of SD8D.

### Oxylipins, not salicylic acid, are responsive to shifts in GlcCer metabolism in *P. patens*

We observed increased accumulation of the cyclopentenones Δ^4^-dn-OPDA and tn-OPDA in GlcCer-deficient mutants, and these accumulations were exacerbated by cold stress treatment in *Ppgcs*. Additionally, RNAseq analysis revealed up-regulation of transcripts associated with oxylipin metabolism in cold-stressed *gcs*. In contrast, SA accumulation was unaffected in the same conditions. In *P. patens* and in the model liverwort *Marchantia polymorpha*, there is accumulating evidence of oxylipins functioning as prominent stress-responsive phytohormones. We speculate that some aspects of the growth phenotypes of GlcCer-deficient mutants may be mediated by oxylipin phytohormones, rather than being direct effects of modified sphingolipid metabolism. Mutagenesis of both pathways will be necessary to understand the distinct effects of sphingolipids vs. oxylipins.

## Acknowledgements

We are grateful to Dr. Jasmin Gömann for generating the initial *gcs* and *sd4d* mutants, and for her intellectual contributions to this project. We would also like to thank Tarek Morsi for IT support for the development of the Cellpose pipeline. We are grateful to Sabine Freitag and Andrea Nickel for technical assistance, and Dr. Ellen Hornung for technical advice throughout this project. We thank Dr. Moritz Klein for advice selecting marker genes for oxylipin metabolism used for interpreting RNAseq data. IF and TH acknowledge funding from the Deutsche Forschungsgemeinschaft (DFG; Priority Programme “MAdLand – Molecular Adaptation to Land: Plant Evolution to Change” SPP 2237: FE 446/14-1, HA 10307/1-1, INST 186/822-1 and INST 186/1167-1).

## Author Contributions

IF and TMH planned and designed the research, and acquired research funding. PD, SB, MR, and DOS performed the experiments. All authors contributed to data analysis. PD, CH, IF, and TMH contributed to method development. TMH wrote the original draft. All authors reviewed the manuscript.

## Data Availability

All data is supplied in the manuscript supplements, and further measurements and metadata are available from the authors upon request.

## Supplemental Data

**Supplemental Figure 1:** Localization of eYFP-PpSD8D transiently expressed in a gametophore phyllid cell following particle bombardment.

**Supplemental Figure 2:** Sphingolipid profile of wt, *sd4d*, *gcs*, and three *sd4d gcs* alleles as gametophores, measured via UPLC-nano-ESI-MS/MS.

**Supplemental Figure 3:** Sphingolipid profile of WT, *sd4d*, *gcs*, and five *sd8d* alleles as gametophores, measured via UPLC-nano-ESI-MS/MS.

**Supplemental Figure 4:** Sphingolipid profile of WT, *sd4d*, two *sd8d* alleles, three *sd4d sd8d* alleles, *gcs*, and *sd4d gcs* as gametophores measured via UPLC-nano-ESI-MS/MS.

**Supplemental Figure 5**: Differentially expressed genes by Pfam and GO term.

**Supplemental Figure 6:** Free ceramides summed according to LCB component, based on UPLC-nano-ESI-MS/MS.

**Supplemental Table S1:** Mutant genotypes used in the study.

**Supplemental Table S2:** Primers used throughout the study.

**Supplemental Table S3:** Sphingolipid profile supporting Figure 2 (gametophores), with 3 alleles for *sd4d sd8d*.

**Supplemental Table S4**: Sphingolipid profile supporting Figure 3 (protonema), with 3 alleles for *sd4d sd8d*.

**Supplemental Table S5:** Sphingolipid profile of mutant gametophores including 3 alleles for *sd4d gcs*.

**Supplemental Table S6:** Sphingolipid profile of mutant gametophores including 5 alleles for *sd8d*.

**Supplemental Table S7:** Complete RNAseq data and comparison of wild type and *gcs* grown in autumn conditions.

**Supplemental Table S8:** All differentially-expressed genes detected with log2FC>1, FDR<0.05.

**Supplemental Table S9**: Genes used for a targeted search for differential regulation of oxylipin and GlcCer metabolism.

## Supplemental Figures

**Supplemental Figure S1:** Localization of eYFP-PpSD8D transiently expressed in a gametophore phyllid cell following particle bombardment. The same cell is imaged in different focal planes to view the nuclear envelope and cortical endoplasmic reticulum. In the merged images, eYFP-PpSD8D is coloured yellow, and SP-mCerulean-KDEL is coloured cyan. Similar localization was observed in four of four cells images expressing this construct. Images were adjusted for brightness and contrast for visualization. Scale bars represent 10 µm. eYFP: enhanced yellow fluorescent protein, SD8D: SPHINGOLIPID Δ8-DESATURASE, SP: signal peptide.

**Supplemental Figure S2:** Sphingolipid profile of wt, *sd4d*(−1), *gcs*(−3), and three *sd4d gcs* alleles (A, B, i) as gametophores, measured via UPLC-nano-ESI-MS/MS. (A) Free LCB totals (B) Free LCB profile (C) Free ceramide totals (D) hydroxylated ceramide totals (E) non-hydroxylated ceramide totals (F) free ceramide profile (G) OH-Cer:nCer ratio (H) GlcCer total (I) GlcCer profile (J) Hex-GIPC (K) HexNAc-GIPC (L) Hex-Hex-GIPC (M) Hex-HexNAc-GIPC (N) Pent-Hex-Hex-GIPC. All unambiguous and robustly-detected lipid species are shown. The peak areas are corrected for exclusion of the ^13^C isotope and normalized to the total FAMEs content of the sample. Data represent the mean ± SD of three replicates grown on separate plates. Statistical analysis was done using a one-way ANOVA with Tukey’s post-hoc test. Letters indicate significance at p < 0.05. Supported by Supplemental Table 5. WT: wild-type, *sd4d*: sphingolipid Δ4-desaturase, *gcs*: glycosylceramide synthase, R; rhizoid, UPLC-nanoESI-MS/MS: ultra high-performance liquid chromatography with nanoelectrospray ionization and triple quadrupole mass spectrometry, FAME: fatty acid methyl ester, LCB: long-chain base, GIPC: glycosyl inositol phosphorylceramide.

**Supplemental Figure S3:** Sphingolipid profile of WT, *sd4d*(−1), *gcs*(−3), and five *sd8d* alleles (A4, A13, B1, B4, B15) as gametophores, measured via UPLC-nano-ESI-MS/MS. (A) Free LCB totals (B) free LCB profile (C) free ceramide totals (D) hydroxylated ceramide totals (E) non-hydroxylated ceramide totals (F) free ceramide profile (G) OH-Cer:nCer ratio (H) GlcCer total (I) GlcCer profile (J) Hex-GIPC (K) HexNAc-GIPC (L) Hex-Hex-GIPC (M) Hex-HexNAc-GIPC (N) Pent-Hex-Hex-GIPC. All unambiguous and robustly-detected lipid species are shown. The peak areas are corrected for exclusion of the ^13^C isotope and normalized to the total FAMEs content of the sample. Data represent the mean ± SD of three replicates grown on separate plates. Statistical analysis was done using a one-way ANOVA with Tukey’s post-hoc test. Letters indicate significance at p < 0.05. Supported by Supplemental Table 6. WT: wild-type, *sd4d*: sphingolipid Δ4-desaturase, *sd8d*: sphingolipid Δ8-desaturase, *gcs*: glycosylceramide synthase, UPLC-nanoESI-MS/MS: ultra high-performance liquid chromatography with nanoelectrospray ionization and triple quadrupole mass spectrometry, FAME: fatty acid methyl ester, FA: fatty acid, LCB: long-chain base, GIPC: glycosyl inositol phosphorylceramide.

**Supplemental Figure S4:** Sphingolipid profile of WT, *sd4d*(−1), two *sd8d* alleles (A4, B1), three *sd4d sd8d* alleles (18,89,69), *gcs*(−3), and *sd4d gcs* (i11) as gametophores measured via UPLC-nano-ESI-MS/MS. (A) Free LCB totals (B) Free LCB profile (C) Free ceramide totals (D) hydroxylated ceramide totals (E) non-hydroxylated ceramide totals (F) free ceramide profile (G) OH to non-OH ceramide ratio (H) GlcCer total (I) GlcCer profile (J) Hex-GIPC (K) HexNAc-GIPC (L) Hex-Hex-GIPC (M) Hex-HexNAc-GIPC (N) Pent-Hex-Hex-GIPC. All unambiguous and robustly-detected lipid species are shown. The peak areas are corrected for exclusion of the ^13^C isotope and normalized to the total FAMEs content of the sample. Data represent the mean ± SD of three replicates grown on separate plates. Statistical analysis was done using a one-way ANOVA with Tukey’s post-hoc test. Letters indicate significance at p < 0.05. Supported by Supplemental Table 3. WT: wild-type, *sd4d*: sphingolipid Δ4-desaturase, *sd8d*: sphingolipid Δ8-desaturase, *gcs*: glycosylceramide synthase, UPLC-nanoESI-MS/MS: ultra high-performance liquid chromatography with nanoelectrospray ionization and triple quadrupole mass spectrometry, FAME: fatty acid methyl ester, FA: fatty acid, LCB: long-chain base, GIPC: glycosyl inositol phosphorylceramide.

**Supplemental Figure S5**: Differentially expressed genes by Pfam (A) and GO term (B). Supported by Supplemental Table 7. Pfam: protein families, GO: gene ontology, FDR: false discovery rate.

**Supplemental Figure S6:** Free ceramides summed according to LCB component, based on UPLC-nanoESI-MS/MS. Supported by Supplemental Table S3, visualized in main Figure 2 Total amounts of free ceramides containing an 18:0;3 LCB are indicated in white. UPLC-nanoESI-MS/MS: ultra high-performance liquid chromatography with nanoelectrospray ionization and triple quadrupole mass spectrometry, FAME: fatty acid methyl ester, WT: wild type, *sd4d*: sphingolipid Δ4-desaturase, *sd8d*: sphingolipid Δ8-desaturase, *gcs*: glycosylceramide synthase.

## Literature Cited

Andersson MX, Larsson KE, Tjellström H, Liljenberg C, Sandelius AS. 2005. Phosphate-limited oat: The plasma membrane and the tonoplast as major targets for phospholipid-to-glycolipid replacement and stimulation of phospholipases in the plasma membrane. Journal of Biological Chemistry 280: 27578–27586.

Bahammou D, Recorbet G, Mamode Cassim A, Robert F, Balliau T, Van Delft P, Haddad Y, Mongrand S, Fouillen L, Simon-Plas F. 2024. A combined lipidomic and proteomic profiling of Arabidopsis thaliana plasma membrane. Plant Journal 119: 1570–1595.

Berkey R, Bendigeri D, Xiao S. 2012. Sphingolipids and Plant Defense/Disease: The “Death” Connection and Beyond. Frontiers in Plant Science 3: 1–22.

Cahoon EB, Kim P, Xie T, Solis AG, Han G, Gong X, Dunn TM. 2025. Sphingolipid homeostasis: How do cells know when enough is enough? Implications for plant pathogen responses. Plant Physiology 197: 1–11.

Carlson M. 2026. org.At.tair.db: Genome wide annotation for Arabidopsis_. R package version 3.22.0.

Carlson M, Falcon S, Pages H, Li N. GO. db: A set of annotation maps describing the entire Gene Ontology. 3: 10–18129.

Carmona-Salazar L, Cahoon RE, Gasca-Pineda J, González-Solís A, Vera-Estrella R, Treviño V, Cahoon EB, Gavilanes-Ruiz M. 2021. Plasma and vacuolar membrane sphingolipidomes: Composition and insights on the role of main molecular species. Plant Physiology 186: 624–639.

Chen M, Markham J, Cahoon EB. 2012. Sphingolipid Δ8 unsaturation is important for glucosylceramide biosynthesis and low-temperature performance in Arabidopsis. Plant Journal 69: 769–781.

Chen M, Markham J, Dietrich CR, Jaworski JG, Cahoon EB. 2008. Sphingolipid long-chain base hydroxylation is important for growth and regulation of sphingolipid content and composition in Arabidopsis. Plant Cell 20: 1862–1878.

Collonnier C, Epert A, Mara K, Maclot F, Guyon-Debast A, Charlot F, White C, Schaefer DG, Nogué F. 2017. CRISPR-Cas9-mediated efficient directed mutagenesis and RAD 51-dependent and RAD 51-independent gene targeting in the moss *Physcomitrella patens*. Plant Biotechnology Journal 15: 122– 131.

Fu W, Pan Y, Shi Y, Chen J, Gong D, Li Y, Hao G, Han D. 2022. Root Morphogenesis of Arabidopsis thaliana Tuned by Plant Growth-Promoting Streptomyces Isolated From Root-Associated Soil of Artemisia annua. Frontiers in Plant Science 12: 1–11.

Gömann J, Herrfurth C, Zienkiewicz K, Haslam TM, Feussner I. 2021a. Sphingolipid Δ4-desaturation is an important metabolic step for glycosylceramide formation in Physcomitrium patens. Journal of Experimental Botany 72: 5569–5583.

Gömann J, Herrfurth C, Zienkiewicz A, Ischebeck T, Haslam TM, Hornung E, Feussner I. 2021b. Sphingolipid long-chain base hydroxylation influences plant growth and callose deposition in *Physcomitrium patens*. New Phytologist 231: 297–314.

Guo Q, Liu L, Rupasinghe TWT, Roessner U, Barkla BJ. 2022. Salt stress alters membrane lipid content and lipid biosynthesis pathways in the plasma membrane and tonoplast. Plant Physiology 189: 805–826.

Haschke H -P, Kaiser G, Martinoia E, Hammer U, Teucher T, Doene AJ, Heinz E. 1990. Lipid Profiles of Leaf Tonoplasts from Plants with Different CO2-Fixation Mechanisms. Botanica Acta 103: 32–38.

Haslam TM, Feussner I. 2022. Diversity in sphingolipid metabolism across land plants. Journal of Experimental Botany 73: 2785–2798.

Haslam TM, Herrfurth C, Feussner I. 2024. Diverse *INOSITOL PHOSPHORYLCERAMIDE SYNTHASE* mutant alleles of *Physcomitrium patens* offer new insight into complex sphingolipid metabolism. New Phytologist 242: 1189–1205.

Herrfurth C, Feussner I. 2020. Quantitative Jasmonate Profiling Using a High-Throughput UPLC-NanoESI-MS/MS Method BT - Jasmonate in Plant Biology: Methods and Protocols. In: Champion A, Laplaze L, eds. Jasmonate in Plant Biology: Methods and Protocols. New York, NY: Springer US, 169– 187.

Herrfurth C, Liu Y-T, Feussner I. 2021. Targeted Analysis of the Plant Lipidome by UPLC-NanoESI-MS/MS. In: Bartels D, Dörmann P, eds. Plant Lipids: Methods and Protocols. Springer Nature, 135–155.

Ito Y, Esnay N, Platre MP, Wattelet-Boyer V, Noack LC, Fougère L, Menzel W, Claverol S, Fouillen L, Moreau P, et al. 2021. Sphingolipids mediate polar sorting of PIN2 through phosphoinositide consumption at the trans-Golgi network. Nature Communications 12: 4267.

Jiang Z, Zhou X, Tao M, Yuan F, Liu L, Wu F, Wu X, Xiang Y, Niu Y, Liu F, et al. 2019. Plant cell-surface GIPC sphingolipids sense salt to trigger Ca2+ influx. Nature 572: 341–346.

König S, Gömann J, Zienkiewicz A, Zienkiewicz K, Meldau D, Herrfurth C, Feussner I. 2022. Sphingolipid-Induced Programmed Cell Death is a Salicylic Acid and EDS1-Dependent Phenotype in Arabidopsis *Fatty Acid Hydroxylase* (*Fah1, Fah2*) and *Ceramide Synthase* (*Loh2*) Triple Mutants. Plant and Cell Physiology 63: 317–325.

Lenarčič T, Albert I, Böhm H, Hodnik V, Pirc K, Zavec AB, Podobnik M, Pahovnik D, Žagar E, Pruitt R, et al. 2017. Eudicot plant-specific sphingolipids determine host selectivity of microbial NLP cytolysins. Science (New York, N.Y.) 358: 1431–1434.

Levine TP, Wiggins CAR, Munro S. 2000. Inositol Phosphorylceramide Synthase is Located in the Golgi Apparatus of *Saccharomyces cerevisiae*. Molecular biology of the cell 11: 2267–2281.

Liu YC, Vidali L. 2011. Efficient polyethylene glycol (PEG) mediated transformation of the moss *Physcomitrella patens*. Journal of Visualized Experiments: 2–5.

Liu P, Xie T, Wu X, Han G, Gupta SD, Zhang Z, Yue J, Dong F, Gable K, Niranjanakumari S, et al. 2023. Mechanism of sphingolipid homeostasis revealed by structural analysis of Arabidopsis SPT-ORM1 complex. Science Advances 9: 1–12.

Lopez-Obando M, Hoffmann B, Géry C, Guyon-Debast A, Téoulé E, Rameau C, Bonhomme S, Nogué F. 2016. Simple and efficient targeting of multiple genes through CRISPR-Cas9 in *Physcomitrella patens*. G3: Genes, Genomes, Genetics 6: 3647–3653.

Love MI, Huber W, Anders S. 2014. Moderated estimation of fold change and dispersion for RNA-seq data with DESeq2. Genome Biology 15: 1–21.

Magnin-Robert M, Le Bourse D, Markham J, Dorey S, Clément C, Baillieul F, Dhondt-Cordelier S. 2015. Modifications of sphingolipid content affect tolerance to hemibiotrophic and necrotrophic pathogens by modulating plant defense responses in Arabidopsis. Plant Physiol. 169: 2255–2274.

Markham J, Li J, Cahoon EB, Jaworski JG. 2006. Separation and Identification of Major Plant Sphingolipid Classes from Leaves. Journal of Biological Chemistry 281: 22684–22694.

Maronova M, Kalyna M. 2016. Generating Targeted Gene Knockout Lines in *Physcomitrella patens* to Study Evolution of Stress-Responsive Mechanisms. Methods Mol Biol. 1398: 221–234.

Michaelson L V., Zäuner S, Markham JE, Haslam RP, Desikan R, Mugford S, Albrecht S, Warnecke D, Sperling P, Heinz E, et al. 2009. Functional characterization of a higher plant sphingolipid Δ4-desaturase: Defining the role of sphingosine and sphingosine-1-phosphate in arabidopsis. Plant Physiology 149: 487–498.

Mittendorf J, Haslam TM, Herrfurth C, Esnay N. 2025. Identification of INOSITOL PHOSPHORYLCERAMIDE SYNTHASE 2 ( IPCS2 ) as a new rate-limiting component in Arabidopsis pathogen entry control. Plant Journal 122: e70159.

Mohammadzadeh R, Karbalaei M, Soleimanpour S, Mosavat A, Rezaee SA, Ghazvini K, Farsiani H. 2021. Practical Methods for Expression of Recombinant Protein in the Pichia pastoris System. Current Protocols 1: e155.

Molino D, Van Der Giessen E, Gissot L, Hématy K, Marion J, Barthelemy J, Bellec Y, Vernhettes S, Satiat-Jeunemaître B, Galli T, et al. 2014. Inhibition of very long acyl chain sphingolipid synthesis modifies membrane dynamics during plant cytokinesis. Biochimica et Biophysica Acta -Molecular and Cell Biology of Lipids 1841: 1422–1430.

Moore WM, Chan C, Ishikawa T, Rennie EA, Wipf HM-L, Benites V, Kawai-Yamada M, Mortimer JC, Scheller H V. 2021. Reprogramming sphingolipid glycosylation is required for endosymbiont persistence in *Medicago truncatula*. Current Biology 31: 1–12.

Mortimer JC, Scheller HV. 2020. Synthesis and Function of Complex Sphingolipid Glycosylation. Trends in Plant Science 25: 522–524.

Msanne J, Chen M, Luttgeharm KD, Bradley AM, Mays ES, Paper JM, Boyle DL, Cahoon RE, Schrick K, Cahoon EB. 2015. Glucosylceramides are critical for cell-type differentiation and organogenesis, but not for cell viability in Arabidopsis. Plant Journal 84: 188–201.

Mueller SJ, Reski R. 2015. Mitochondrial Dynamics and the ER: The Plant Perspective. Front. Cell Dev. Biol. 3: 78.

Nightingale A, Antunes R, Alpi E, Bursteinas B, Gonzales L, Liu W, Luo J, Qi G, Turner E, Martin M. 2017. The Proteins API: Accessing key integrated protein and genome information. Nucleic Acids Research 45: W539–W544.

Resemann HC, Herrfurth C, Feussner K, Hornung E, Ostendorf AK, Gömann J, Mittag J, van Gessel N, Vries J de, Ludwig-Müller J, et al. 2021. Convergence of sphingolipid desaturation across over 500 million years of plant evolution. Nature Plants 7: 219–232.

Saavedra L, Catarino R, Heinz T, Heilmann I, Bezanilla M, Malhó R. 2015. Phosphatase and tensin homolog is a growth repressor of both rhizoid and gametophore development in the moss Physcomitrella patens. Plant Physiology 169: 2572–2586.

Sears IB, O’Connor J, Rossanese OW, Glick BS. 1998. A versatile set of vectors for constitutive and regulated gene expression in Pichia pastoris. Yeast 14: 783–790.

Slowikowski K. 2026. ggrepel: Automatically Position Non-Overlapping Text Labels with ‘ggplot2’_. R package version 0.9.8.

Steinberger AR, Merino WO, Cahoon RE, Cahoon EB, Lynch D V. 2021. Disruption of long-chain base hydroxylation alters growth and impacts sphingolipid synthesis in *Physcomitrella patens*. Plant Direct 5: 1–19.

Stringer C, Wang T, Michaelos M, Pachitariu M. 2021. Cellpose: a generalist algorithm for cellular segmentation. Nature Methods 18: 100–106.

Team RC. 2026. A Language and Environment for Statistical Computing. R Foundation for Statistical Computing.

Ternes P, Wobbe T, Schwarz M, Albrecht S, Feussner K, Riezman I, Cregg JM, Heinz E, Riezman H, Feussner I, et al. 2011. Two pathways of sphingolipid biosynthesis are separated in the yeast Pichia pastoris. Journal of Biological Chemistry 286: 11401–11414.

Wegner L, Herrfurth C, Feussner I, Ehlers K, Haslam TM. 2025. Complex sphingolipid metabolism impacts cell division and plasmodesmal development in the moss Physcomitrium patens. Plant Physiology 199: 1–19.

Weidner M, Taupp M, Hallam SJ. 2010. Expression of recombinant proteins in the methylotrophic yeast Pichia pastoris. Journal of Visualized Experiments: 1–5.

Wickham H. 2016. ggplot2: Elegant Graphics for Data Analysis. New York: Springer-Verlag.

Xu S, Hu E, Cai Y, Xie Z, Luo X, Zhan L, Tang W, Wang Q, Liu B, Wang R, et al. 2024. Using clusterProfiler to characterize multiomics data. Springer US.

Zienkiewicz A, Gömann J, König S, Herrfurth C, Liu YT, Meldau D, Feussner I. 2020. Disruption of Arabidopsis neutral ceramidases 1 and 2 results in specific sphingolipid imbalances triggering different phytohormone-dependent plant cell death programmes. New Phytologist 226: 170–188.

