## Supplemental Figures for "Glycosylceramide assembly and function in a model bryophyte"

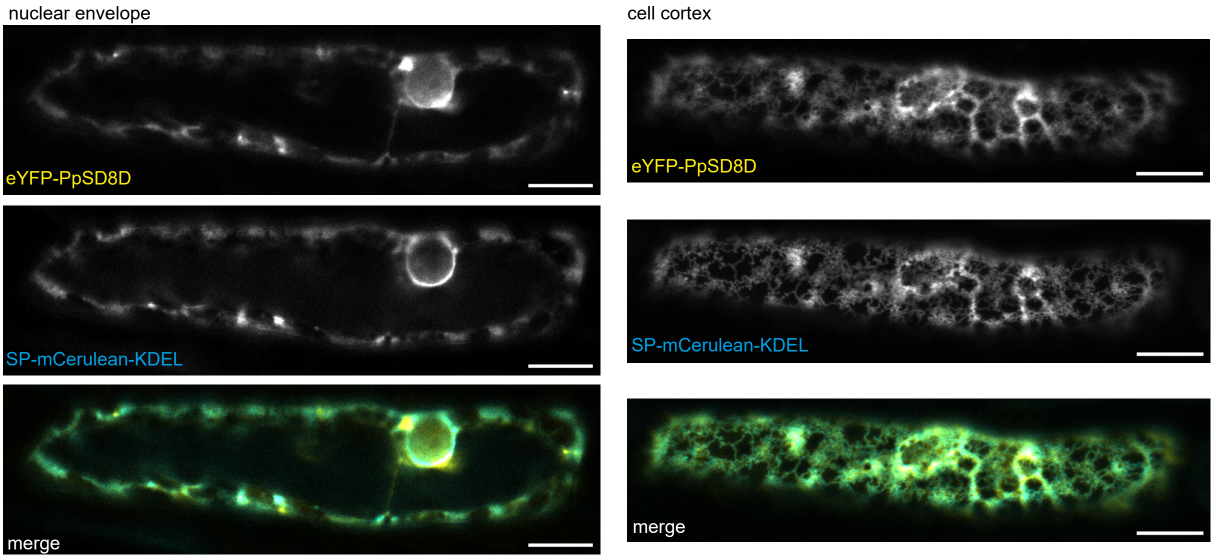


**Supplemental Figure S1**: Localization of eYFP-PpSD8D transiently expressed in a gametophore phyllid cell following particle bombardment. The same cell is imaged in different focal planes to view the nuclear envelope and cortical endoplasmic reticulum. In the merged images, eYFP-PpSD8D is coloured yellow, and SP-mCerulean-KDEL is coloured cyan. Similar localization was observed in four of four cells imagesd expressing this construct. Images were adjusted for brightness and contrast for visualization. Scale bars represent 10 µm. eYFP: enhanced yellow fluorescent protein, SD8D: SPHINGOLIPID DELTA-8 DESATURASE, SP: signal peptide.

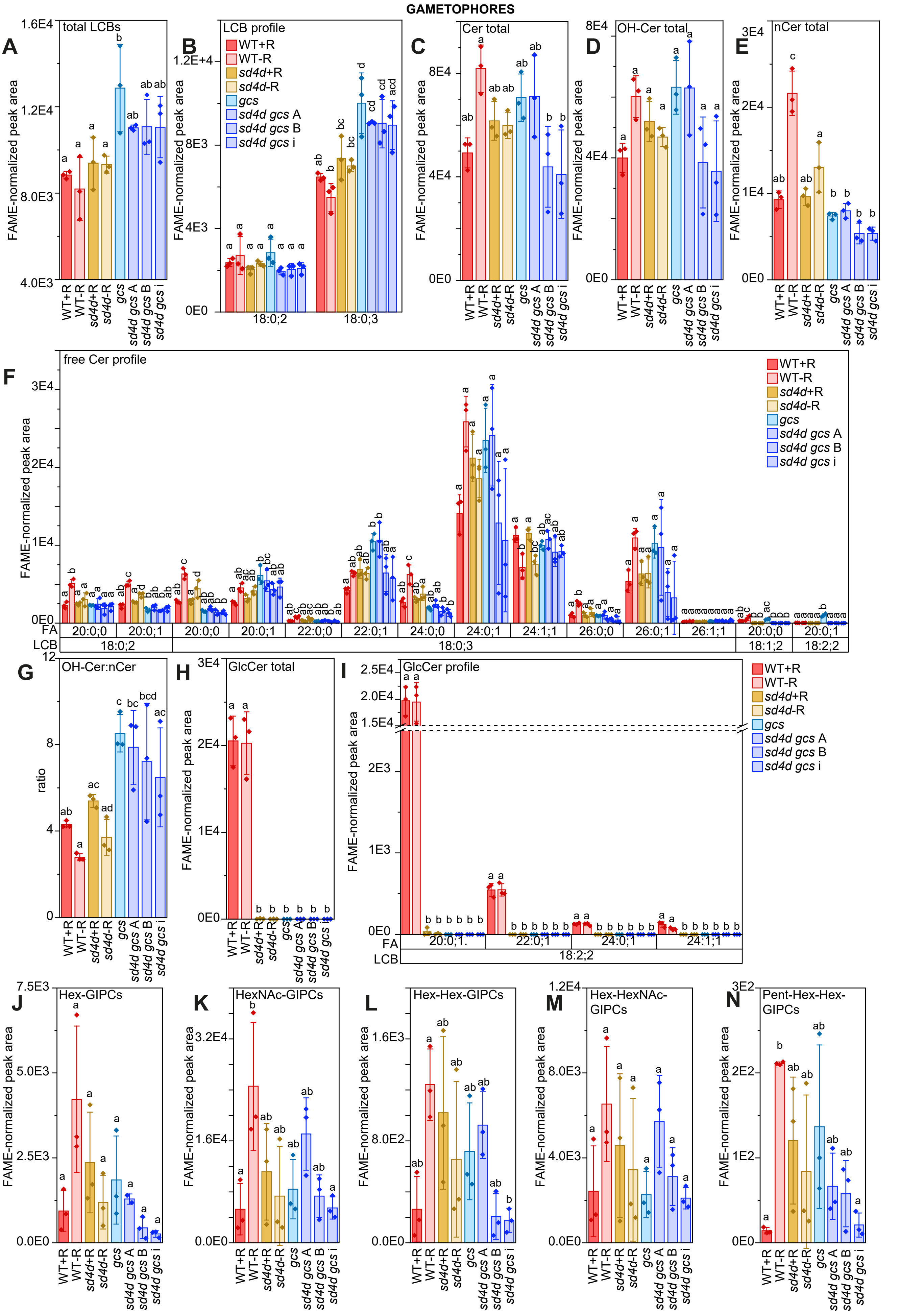


**Supplemental Figure S2:** Sphingolipid profile of wt, *sd4d*(-1), *gcs*(-3), and three *sd4d gcs* alleles (A, B, i) as gametophores, with +/- rhizoid controls, measured via UPLC-nano-ESI-MS/MS. (A) Free LCB totals (B) Free LCB profile (C) Free ceramide totals (D) hydroxylated ceramide totals (E) non-hydroxylated ceramide totals (F) free ceramide profile (G) OH-Cer:nCer ratio (H) GlcCer total (I) GlcCer profile (J) Hex-GIPC (K) HexNAc-GIPC (L) Hex-Hex-GIPC (M) Hex-HexNAc-GIPC (N) Pent-Hex-Hex-GIPC. All unambiguous and robustly-detected lipid species are shown. The peak areas are corrected for exclusion of the C13 isotope and normalized to the total FAMEs content of the sample. Data represent the mean ± SD of three replicates grown on separate plates. Statistical analysis was done using a one-way ANOVA with Tukey's post-hoc test. Letters indicate significance at $p < 0.05$. Supported by Supplemental Table 5. WT: wild-type, *sd4d*: sphingolipid delta-4 desaturase, *gcs*: glycosylceramide synthase, R; rhizoid, UPLC-nanoESI-MS/MS: ultra high-performance liquid chromatography with nanoelectrospray ionization and triple quadrupole mass spectrometry, FAME: fatty acid methyl ester, FA: fatty acid, LCB: long-chain base, GIPC: glycosyl inositol phosphorylceramide.

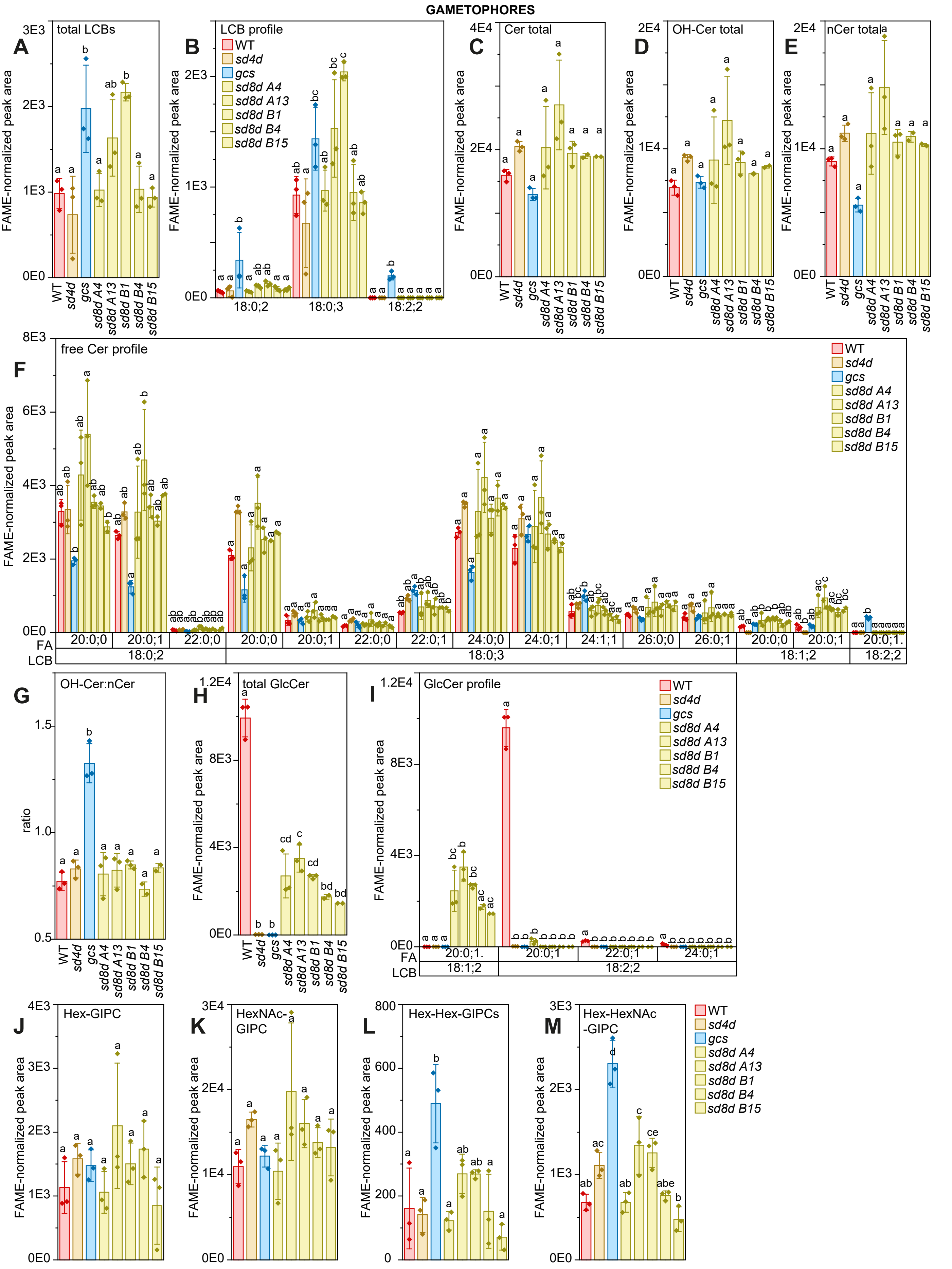


**Supplemental Figure S3:** Sphingolipid profile of WT, *sd4d*(-1), *gcs*(-3), and five *sd8d* alleles (A4, A13, B1, B4, B15) as gametophores, measured via UPLC-nano-ESI-MS/MS. (A) Free LCB totals (B) free LCB profile (C) free ceramide totals (D) hydroxylated ceramide totals (E) non-hydroxylated ceramide totals (F) free ceramide profile (G) OH-Cer:nCer ratio (H) GlcCer total (I) GlcCer profile (J) Hex-GIPC (K) HexNAc-GIPC (L) Hex-Hex-GIPC (M) Hex-HexNAc-GIPC (N) Pent-Hex-Hex-GIPC. All unambiguous and robustly-detected lipid species are shown. The peak areas are corrected for exclusion of the $^{13}C$ isotope and normalized to the total FAMEs content of the sample. Data represent the mean ± SD of three replicates grown on separate plates. Statistical analysis was done using a one-way ANOVA with Tukey's post-hoc test. Letters indicate significance at $p < 0.05$. Supported by Supplemental Table 6. WT: wild-type, *sd4d*: sphingolipid Δ4-desaturase, *sd8d*: sphingolipid Δ8-desaturase, *gcs*: glycosylceramide synthase, UPLC-nanoESI-MS/MS: ultra high-performance liquid chromatography with nanoelectrospray ionization and triple quadrupole mass spectrometry, FAME: fatty acid methyl ester, FA: fatty acid, LCB: long-chain base, GIPC: glycosyl inositol phosphorylceramide.

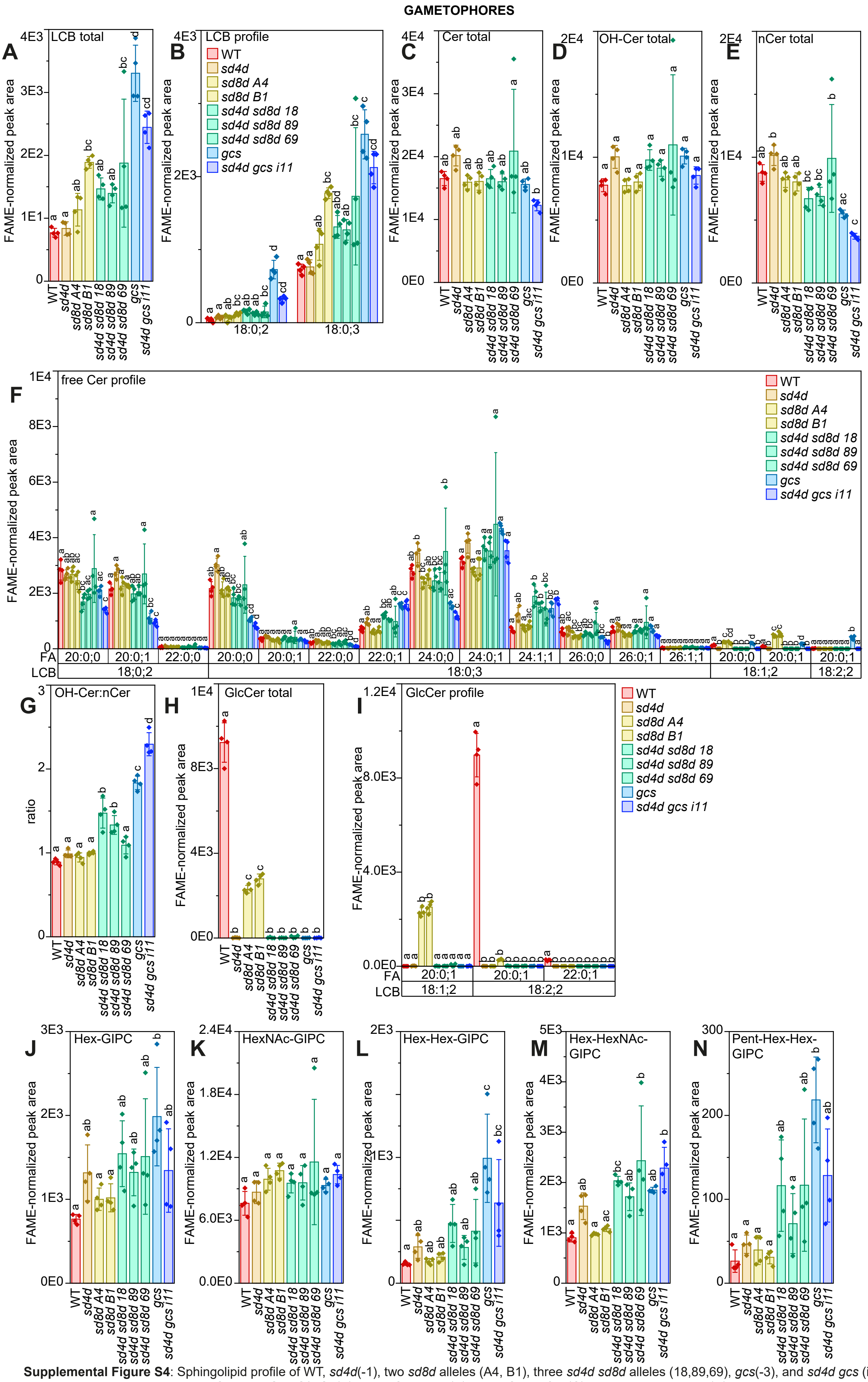


**Supplemental Figure S4**: Sphingolipid profile of WT, *sd4d*(-1), two *sd8d* alleles (A4, B1), three *sd4d sd8d* alleles (18,89,69), *gcs*(-3), and *sd4d gcs* (i11) as gametophores measured via UPLC-nano-ESI-MS/MS. (A) Free LCB totals (B) Free LCB profile (C) Free ceramide totals (D) hydroxylated ceramide totals (E) non-hydroxylated ceramide totals (F) free ceramide profile (G) OH to non-OH ceramide ratio (H) GlcCer total (I) GlcCer profile (J) Hex-GIPC (K) HexNAc-GIPC (L) Hex-Hex-GIPC (M) Hex-HexNAc-GIPC (N) Pent-Hex-Hex-GIPC. All unambiguous and robustly-detected lipid species are shown. The peak areas are corrected for exclusion of the $^{13}$C isotope and normalized to the total FAMEs content of the sample. Data represent the mean ± SD of three replicates grown on separate plates. Statistical analysis was done using a one-way ANOVA with Tukey's post-hoc test. Letters indicate significance at $p < 0.05$. Supported by Supplemental Table 3. WT: wild-type, *sd4d*: sphingolipid Δ4-desaturase, *sd8d*: sphingolipid Δ8-desaturase, *gcs*: glycosylceramide synthase, UPLC-nanoE-SI-MS/MS: ultra high-performance liquid chromatography with nanoelectrospray ionization and triple quadrupole mass spectrometry, FAME: fatty acid methyl ester, FA: fatty acid, LCB: long-chain base, GIPC: glycosyl inositol phosphorylceramide.

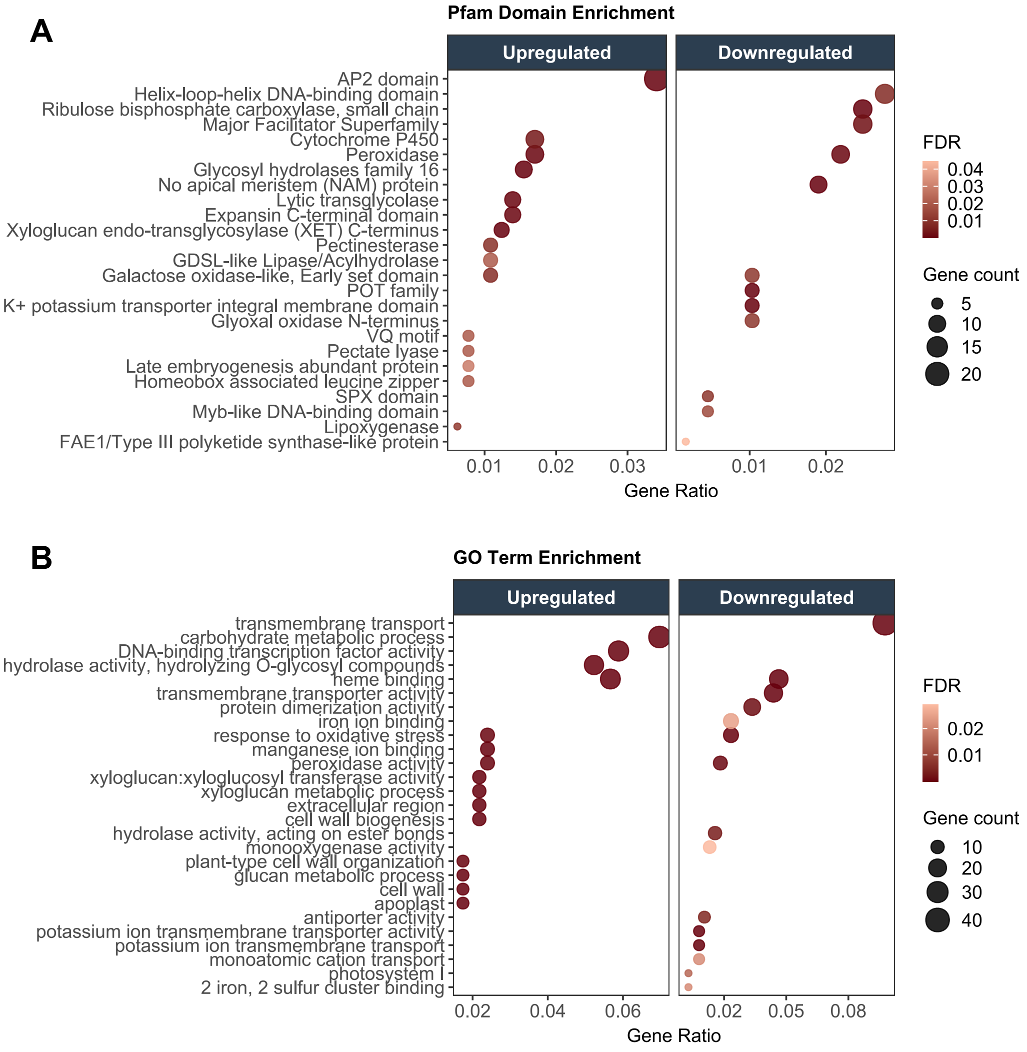


**Supplemental Figure S5:** Differentially expressed genes by Pfam (A) and GO term (B). Supported by Supplemental Table 7. Pfam: protein families, GO: gene ontology, FDR: false discovery rate.

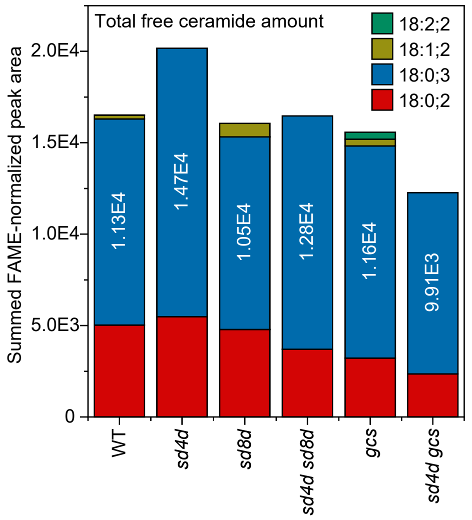


**Supplemental Figure S6**: Free ceramides summed according to LCB component, based on UPLC-nanoESI-MS/MS. Supported by Supplemental Table S3, visualized in main Figure 2. Total amounts of free ceramides containing an 18:0;3 LCB are indicated in white. UPLC-nanoESI-MS/MS: ultra high-performance liquid chromatography with nanoelectrospray ionization and triple quadrupole mass spectrometry, FAME: fatty acid methyl ester, WT: wild type, *sd4d*: sphingolipid Δ4-desaturase, *sd8d*: sphingolipid Δ8-desaturase, *gcs*: glycosylceramide synthase.
